# Trapping a somatic endogenous retrovirus into a germline piRNA cluster immunizes the germline against further invasion

**DOI:** 10.1101/510016

**Authors:** Céline Duc, Marianne Yoth, Silke Jensen, Nolwenn Mouniée, Casey M. Bergman, Chantal Vaury, Emilie Brasset

## Abstract

**Background:** For species survival, the germline must faithfully transmit genetic information to the progeny. Transposable elements (TEs), which are major components of eukaryotic genomes, constitute a significant threat to genome stability due to their mobility. In the metazoan germline, their mobilization is limited by a class of small RNAs that are called PIWI-interacting RNAs (piRNAs) and are produced by dedicated genomic loci called piRNA clusters. Although the piRNA pathway is an adaptive genomic immunity system, it remains unclear how the germline is protected from a new transposon invasion. To address this question, we used *Drosophila melanogaster* lines harboring a deletion within *flamenco*, a major piRNA cluster that is specifically expressed in somatic follicular cells. This deletion leads to derepression of the retrotransposon *ZAM* in the somatic follicular cells and subsequent germline genome invasion.

**Results:** In this mutant line that express *ZAM* in somatic follicular cells, we identified *de novo* production of sense and antisense *ZAM*-derived piRNAs that displayed a germinal molecular signature. These piRNAs originated from a new *ZAM* insertion into a germline dual-strand piRNA cluster and silenced *ZAM* expression specifically in germ cells. Finally, we found that *ZAM* trapping in a germinal piRNA cluster is a frequent event that occurs early during the isolation of the mutant line.

**Conclusions:** Transposons can hijack the host developmental process to propagate whenever their silencing is lost. Here, we show that the germline can protect itself by trapping invading somatic-specific TEs into germline piRNA clusters. This is the first demonstration of “auto-immunization” of the germline endangered by mobilization of a surrounding somatic TE.

## Background

Germ cells are the only cell type within multicellular organisms that can transfer genetic and epigenetic material to the offspring. Due to their capacity to move, transposable elements (TEs), a major component of eukaryotic genomes, constitute a significant threat to the germline genome integrity [1–3]. Indeed, their mobilization could lead to gene disruption or chromosomal rearrangements. To limit TE mobilization in the germline, a class of small RNAs of 23 to 29 nucleotides (nt) in length, called PIWI-interacting RNA (piRNAs), are expressed in the reproductive tissue and silence TE activity via homology-dependent mechanisms [4–7].

The piRNA pathway has been extensively studied in the *Drosophila melanogaster* ovary that comprises about sixteen ovarioles, each of which contains a succession of follicles composed of germline and somatic follicular cells [8]. In *D. melanogaster*, piRNAs are encoded by dedicated genomic loci that are called piRNA clusters [9]. These clusters are composed of full length or truncated TEs that define the repertoire of elements that are recognized and silenced by the piRNA machinery. Two classes of piRNA clusters have been defined on the basis of their transcriptional properties: (i) unidirectional or uni-strand, and (ii) bidirectional or dual-strand piRNA clusters [9]. Unidirectional clusters are expressed predominantly in somatic follicular cells of ovaries, while bidirectional clusters are transcribed in germline cells. Therefore, TEs are silenced in both cell types by piRNAs *via* different mechanisms [10,11]. Transcription of piRNA clusters produces long piRNA precursors that are diced into piRNAs. In germline cells, these piRNAs are loaded on the Piwi protein to form a complex that triggers TE transcriptional silencing [12]. In addition to Piwi, two other PIWI-family proteins, Aub and Ago3, participate in the post-transcriptional control of TEs. They act to amplify the piRNA pool by a mechanism called the ping-pong cycle [9]. Moreover, Aub- and Ago3-bound piRNAs are deposited in the embryo to ensure the re-initiation of piRNA clusters and efficient TE control in the offspring germline [13–15]. In somatic follicular cells, whose genome does not contribute to the next generation but which could be the origin of transposon invasion, a simplified version of the piRNA pathway is active because only the Piwi protein is expressed [16,17]. The tissue-specific expression of piRNA clusters, which contain different TE sequences, suggests a tissue-specific regulation of certain classes of elements. For instance, *flamenco* is the best characterized piRNA cluster predominantly expressed in somatic follicular cells. The *flamenco* locus is a uni-strand cluster that extends over more than 180 kb and is located in the pericentromeric heterochomatin of *D. melanogaster* X chromosome [18–20]. Most TEs inserted in *flamenco* belong to the long terminal repeat (LTR) group of retrotransposons and are oriented opposite to the cluster transcription direction. Across the entire spectrum of transposons described in *flamenco*, maternally deposited piRNAs targeting some TEs, such as *ZAM* or *gypsy*, are underrepresented in the embryonic piRNA pool [16]. This suggests that piRNAs matching these TEs are not produced by any germline piRNA cluster and that they originate from the main somatic piRNA cluster, *flamenco*. Thus, these TEs should be exclusively silenced in somatic follicular cells. In the absence of efficient silencing of these TEs in somatic follicular cells, the oocyte genome is exposed to internal threats. Indeed, when the silencing of *ZAM* or *gypsy* is released in somatic follicular cells, these (and potentially other) retrovirus-like TEs can infect germline cells [21,22]. Therefore, the stability of the germline genome requires efficient silencing of TEs also in somatic follicular cells.

The piRNA pathway has often been compared to an adaptive immune system, because it conveys the memory of previous transposon invasions by storing TE sequence information within piRNA clusters [16]. This model leads to several major questions. Particularly, it is not known whether some TE classes are regulated only in specific tissues and whether and how germ cells can counteract TE invasion from the surrounding somatic follicular cells. To gain insights into these issues, we used *D. melanogaster* lines in which *ZAM* expression is either silenced (i.e., “stable”, w^IR6^ line) or derepressed (i.e., “unstable”, RevI-H2 also named RevI in [20]). The RevI-H2 line was derived from the w^IR6^ line after P-mediated mutagenesis [23,24] and displays a large deletion of the proximal *(i.e*. the region closest to the centromere) part of *flamenco* corresponding to the region containing its only *ZAM* insertion [25]. This suggests a tight correlation between the presence of *ZAM* in the *flamenco* locus and the repression of all functional genomic copies of *ZAM* in the somatic follicular cells [25].

Here, we found that in the w^IR6^ ovaries, *ZAM* was silenced only in follicular cells with an absence of a germline-specific silencing mechanism. Conversely, in the RevI-H2 line, ZAM was derepressed in somatic follicular cells and silenced in the germline following its rapid trapping into a germline piRNA cluster. This represents an efficient mechanism of protection against TE invasion from the surrounding somatic tissues.

## Results

### *ZAM* is silenced in a tissue-specific manner

Previous studies have reported that distinct tissue-specific piRNA populations are expressed in the germline and in somatic follicular cells [16]. This suggests a tissue-specific repression of TEs. Here, we used *ZAM* to monitor the germline capacity to repress TEs for which no germline piRNA is produced. *ZAM* is a prototypic somatic TE [16,26] and *ZAM*-derived piRNAs are highly depleted in the early embryonic piRNA population that mirrors the germline piRNA population [16]. To monitor *ZAM* repression, we generated a sensor transgene that expresses the *GFP* reporter gene under the control of an inducible Upstream Activation Sequence promoter (UASp) and harbors a *ZAM* fragment in its 3’UTR (p*GFP-ZAM*) (Fig. 1A). Transgene expression analysis in both somatic and germline cells using the *actin*-Gal4 driver showed that p*GFP-ZAM* was completely silenced in somatic cells (Fig. 1B). This indicated that *ZAM*-derived piRNAs, which are produced by *flamenco* in these cells, targeted the transgene and efficiently guided its silencing. Conversely, in germline cells its expression was not inhibited, as shown by the strong GFP signal (Fig. 1B). This showed that *ZAM* silencing is specific to somatic follicular cells suggesting that it is mediated by the somatic *flamenco* cluster, as shown by genetic evidence [20,27], and that there are no *ZAM*-derived piRNAs from any germline piRNA cluster.

**Figure 1.**
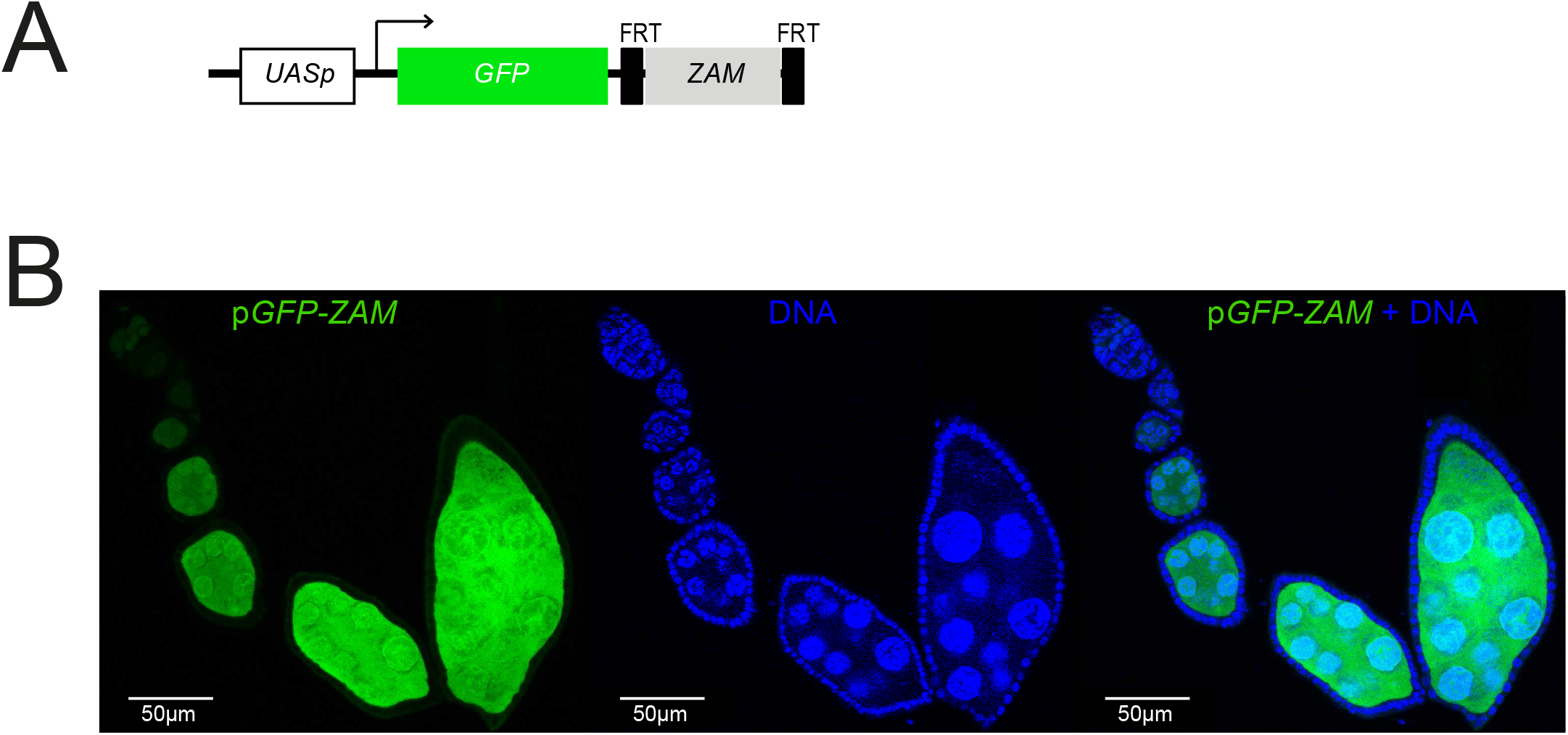
The *ZAM* sensor transgene is not repressed in the germline of *D. melanogaster* ovaries. **a** Structure of the p*GFP-ZAM* sensor transgene. The UASp promoter contains the Gal4 target sequence upstream of the *GFP* reporter gene fused to 467bp of the *ZAM env* gene (light grey box, sense orientation). The *ZAM* sequence is flanked by two FRT sites. The arrow indicates the transcription initiation site. **b** Confocal images of ovarioles after GFP (green, left) and DNA (blue, middle) staining. Ovarioles were from the progeny of a cross between w^1118^ females and males harboring the p*GFP-ZAM* transgene driven by the *actin*-Gal4 driver. Merged images for GFP and DNA labeling are displayed on the right.

### *ZAM*-derived piRNAs are produced in the germline in response to follicular cell instability

*ZAM* silencing release in somatic follicular cells could expose the oocyte genome to internal threats arising from the surrounding follicular cells. To analyze how the germline may protect itself against TE mobilization from the surrounding follicular cells, we used RevI-H2 flies harboring a deletion in the proximal part of *flamenco* [20] that eliminates the region in which *ZAM* is inserted [25] (Fig.S1A), but does not affect germline development. In contrast, as the *flamenco* piRNA cluster is the main source of piRNAs (78%) produced in somatic follicular cells (Fig. 2A), other mutations affecting *flamenco* expression, such as *flamKG* and *flamBG*, lead to disruption of piRNA production, but also to impairment of ovarian germline stem cell differentiation and division, thus preventing further analysis of how the germline might respond to any TE mobilization initiated in the surrounding follicular cells [27]. In addition, the close relationship between the parental w^IR6^ and derived RevI-H2 allowed us to closely control for the genetic background.

**Figure 2.**
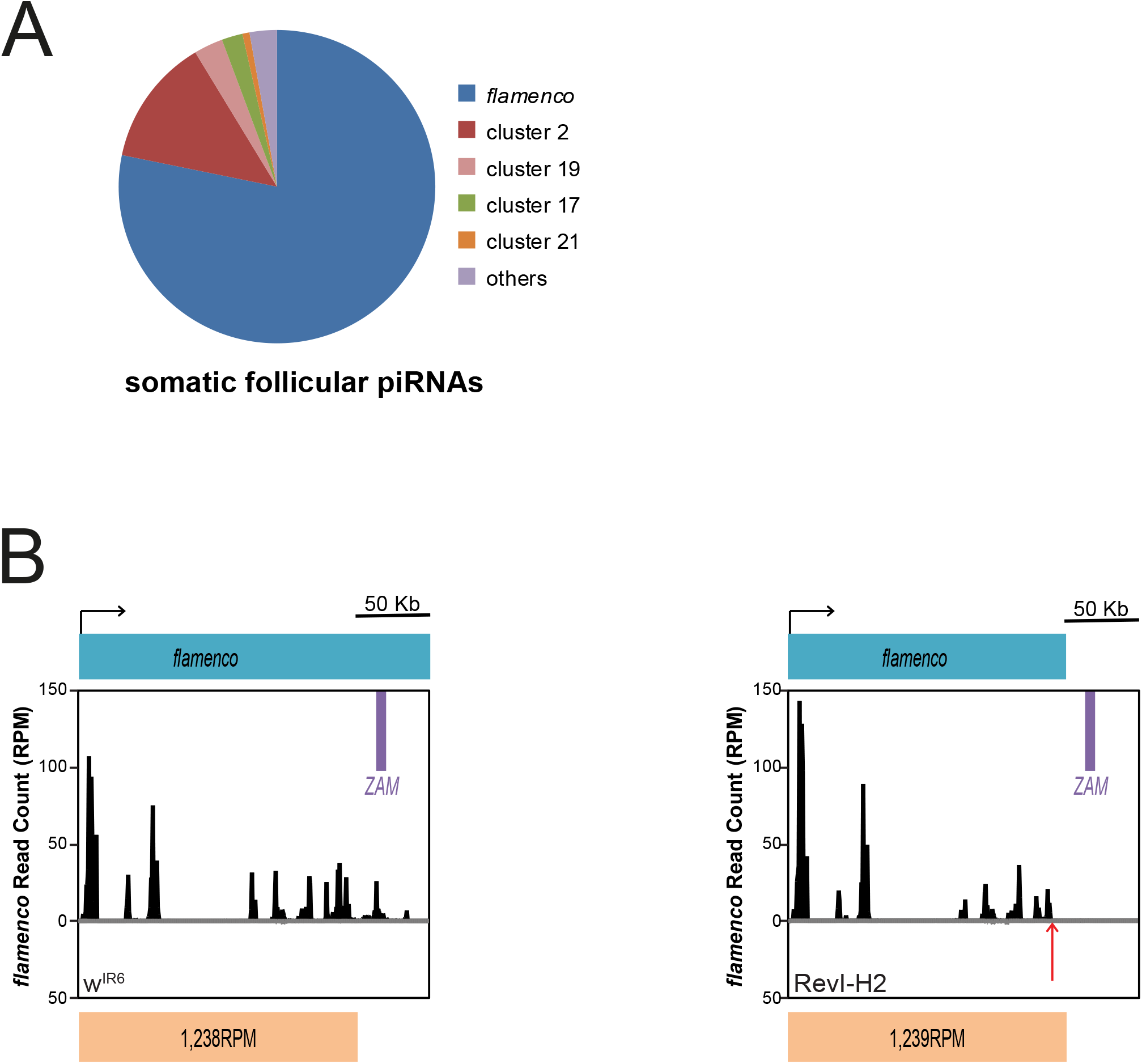
Deletion of some TE fragments in *flamenco* does not impair the global piRNA production from this piRNA cluster. **a** Pie chart showing the proportion of unique piRNAs that map to each of the 142 piRNA clusters in ovarian somatic sheath cells (no mismatch allowed, piRNA clusters defined as in [9]). **b** Density profile of unique piRNAs from the w^IR6^ (left) and RevI-H2 (right) lines that map to the *flamenco* piRNA cluster. Sense and antisense reads are presented in black and grey, respectively. Almost no antisense reads map to the *flamenco* piRNA cluster. *ZAM* location in *flamenco* is displayed by a purple box. The *flamenco* deletion distal break-point in RevI-H2 [25] (Fig S1B) is indicated by a red arrow and the sense of transcription by a black arrow. The count of piRNA reads per million (RPM) mapping the non-deleted region of *flamenco*, indicated below, does not differ between w^IR6^ and RevI-H2.

To determine whether the *flamenco* deletion in RevI-H2 was associated with changes in piRNA production at this locus, we sequenced and compared ovarian small RNAs from the RevI-H2 line and the parental w^IR6^ line. This highlighted the complete loss of piRNAs produced at the deleted locus in RevI-H2 samples compared with the w^IR6^ control line (Fig. 2B). Conversely, the global production of piRNAs uniquely mapping to the *flamenco* locus upstream of the deletion was not affected by the deletion (1,238 and 1,239 Reads Per Million for the RevI-H2 and w^IR6^ samples, respectively) (Fig. 2B and S1B).

As expected from earlier studies, in the w^IR6^ control line, 88% of *ZAM*-derived piRNAs mapped to piRNA clusters [9] (without mismatch) and 86% of them mapped the *flamenco* locus (Fig. 3A). Detailed analysis showed that piRNAs were predominantly antisense to the *ZAM* sequence (Fig. 3B), in agreement with *ZAM* insertion in the antisense orientation relative to *flamenco* transcription orientation (Fig S1A). [25]. Moreover, 90% of *ZAM*-derived piRNAs displayed a uridine bias at the 5’ end, a feature of mature primary piRNAs (Fig. 3C). As *ZAM* is absent from the RevI-H2 *flamenco* locus and is derepressed in somatic follicular cells of RevI-H2 ovaries [20], we hypothesized that production of *ZAM*-derived piRNAs would be abolished in RevI-H2 ovaries. However, sequencing of ovarian small RNAs revealed that antisense *ZAM*-derived piRNAs were considerably increased (three times) in RevI-H2 ovaries compared with w^IR6^ ovaries (Fig. 3D). Moreover, many more *ZAM*-derived sense piRNAs were produced in RevI-H2 than in w^IR6^ ovaries (Fig. 3E). To identify the cellular origin of these *ZAM*-derived piRNAs, we performed a nucleotide profile analysis. We identified a bias for uracil at the first position (1U) and for adenine at the tenth position (10A) (Fig. 3F). This is a typical feature of piRNAs generated by the ping-pong amplification mechanism that occurs exclusively in germline cells. We then checked the ping-pong signature *(i.e*., a 10-nucleotide overlap between sense and antisense pairs of *ZAM*-derived piRNAs) [9] and found a significant enrichment for this signature in the RevI-H2 line, but not in the parental w^IR6^ line (Fig. 3G). Moreover, in RevI-H2 samples, 34% of the *ZAM*-derived piRNAs possessed ping-pong partners (PPP), *i.e*. piRNAs which present a 10-nt 5’-overlap between sense and antisense *ZAM*-derived piRNAs (Fig. 3H). In addition, they harbored the typical 10A and 1U bias (Fig. 3I, Fig. S2A-B). This abundant production of sense and antisense *ZAM*-derived piRNAs and the ping-pong signature enrichment were similar to the results obtained for piRNAs derived from *Burdock*, a typical target of the germline piRNA pathway (Fig. S2C-H). Altogether, these findings strongly suggested a germinal origin of the *ZAM*-derived piRNAs produced in the RevI-H2 line.

**Figure 3.**
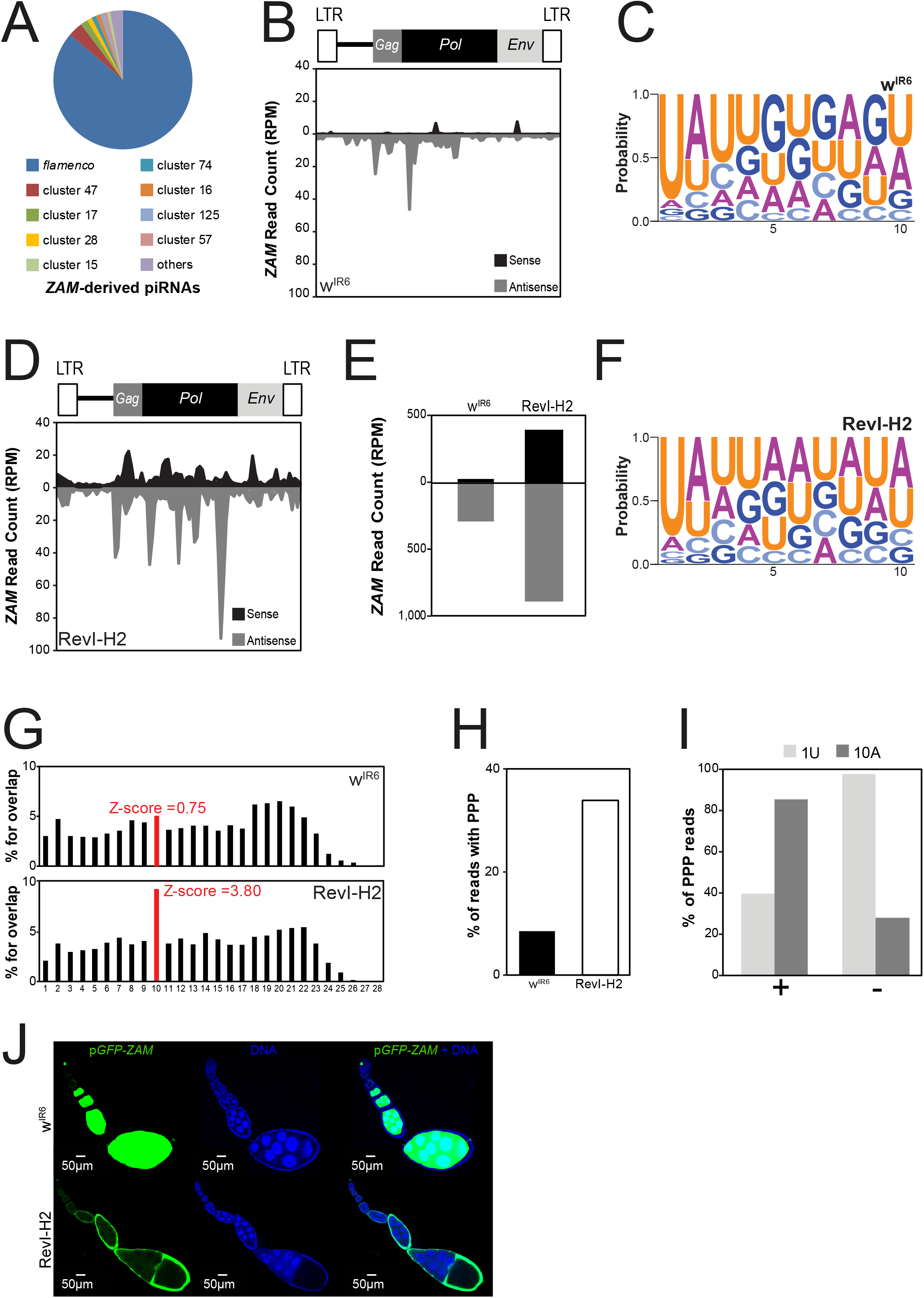
*De novo* production of functional *ZAM*-derived piRNAs in the germline of the RevI-H2 line. **a** Pie chart summarizing the proportion of *ZAM*-derived piRNAs (allowing up to 3 mismatches) that map to the 142 piRNA clusters in w^IR6^ (no mismatch allowed, piRNA clusters defined as in [9]). **b** Density profile of *ZAM*-derived piRNAs along the 8.4kb *ZAM* sequence in w^IR6^ ovaries (allowing up to 3 mismatches). Sense and antisense reads are represented in black and grey, respectively. *ZAM* organization is displayed above the profile. LTR, long terminal repeats. **c** Logo of nucleotide bias for the first ten positions in *ZAM*-derived piRNAs produced in w^IR6^ ovaries. The nucleotide height represents its relative frequency at that position. **d** Density profile of *ZAM*-derived piRNAs along the *ZAM* sequence produced in RevI-H2 ovaries (allowing up to 3 mismatches). Sense and antisense reads are represented in black and grey, respectively. **e** Bar diagram showing the total amount of *ZAM*-derived piRNAs produced in w^IR6^ and RevI-H2 ovaries, quantified from the profiles shown in *b* and *c*, respectively. **f** Logo of nucleotide bias for the first ten positions of *ZAM*-derived piRNAs produced in RevI-H2 ovaries. **g** Histogram showing the percentage of 5’-overlap between sense and antisense *ZAM*-derived piRNAs (23-29nt) in w^IR6^ (top) and RevI-H2 (bottom) ovaries. The proportion of 10-nt overlapping pairs is in red and the Z-score is indicated. **h** Bar diagram indicating the percentage of *ZAM*-derived piRNAs with ping-pong partners (PPP) in the w^IR6^ and RevI-H2 lines. **i** Analysis of nucleotide bias for sense (+) and antisense (−) *ZAM*-derived piRNAs with PPP in RevI-H2 ovaries. The percentage of PPP with a uridine at position 1 (1U) and with an adenosine at position 10 (10A) is shown. **j** Confocal images of ovarioles after GFP (green, left panels) and DNA (blue, middle panels) staining. Ovarioles were from the progeny of a cross between w^IR6^ or RevI-H2 females and males carrying the p*GFP-ZAM* sensor transgene driven by *actin*-Gal4. Right panels, merged images of GFP and DNA labeling.

Aub and Ago3, the two main proteins involved in piRNA production through the ping-pong mechanism, were expressed only in the germline in both w^IR6^ and RevI-H2 ovaries (Fig. S3A-B). This excluded a ping-pong-mediated ectopic production of *ZAM*-derived piRNAs in somatic cells of RevI-H2 ovaries. Moreover, we found that these new *ZAM*-derived piRNAs in RevI-H2 were maternally deposited in early embryos (Fig. S3C-D) and possessed the same characteristics as those produced in adult ovaries (Fig. S3E-G). Taken together, our data strongly suggested that these *ZAM*-derived piRNAs were produced in the germline of RevI-H2 ovaries. This is intriguing because *ZAM* has been classified as a somatic TE, only expressed in somatic cells [16,20].

To monitor the silencing potential of *ZAM*-derived piRNAs produced in the germline of the RevI-H2 ovaries, we followed the GFP expression of the p*GFP-ZAM* sensor transgene in the presence of the *actin*-Gal4 driver. In w^IR6^ control ovaries, the transgene was completely silenced in somatic cells and strongly expressed in germline cells (Fig. 3J) as observed for w^1118^ (Fig. 1B). Conversely, in RevI-H2 ovaries, the transgene was silenced in the germline and strongly expressed in somatic cells. When the *ZAM* sequence was excised upon recombination between the flanking FRTs giving rise to a p*GFP* transgene lacking targets for *ZAM* piRNAs, *GFP* is strongly expressed in RevI-H2 and w^IR6^ somatic and germline cells indicating that the *ZAM* fragment in the fusion transcript is responsible for *GFP* repression (Fig. S3H). To confirm that the p*GFP-ZAM* transgene silencing is piRNAs mediated, we knocked down the germline expression of Aub or AGO3 and monitored GFP expression. We showed that the p*GFP-ZAM* transgene is strongly expressed in the germline of both *Aub*-KD and AGO3-KD (Fig. S3I-J), confirming that the transgene is silenced by a piRNA-mediated mechanism. These results indicated that RevI-H2 germline cells produce *ZAM*-derived piRNAs that efficiently guide sensor silencing. Conversely, GFP is strongly expressed in RevI-H2 somatic follicular cells that do not produce ZAM-derived piRNAs due to the deletion of the proximal part of *flamenco*.

Taken together, we concluded that in RevI-H2 ovaries, functional *ZAM*-derived piRNAs are produced in the germline from a new *ZAM* insertion somewhere outside the deleted region of the *flamenco* cluster.

### *ZAM* transposed into a pre-existing germline piRNA cluster

*ZAM*-derived piRNA production in the RevI-H2 line could be explained by insertion of a new copy of *ZAM* into a pre-existing germline piRNA cluster or by the *de novo* creation of a piRNA cluster in the germline induced by a new *ZAM* insertion. To discriminate between these hypotheses, we studied the activity of this putative piRNA cluster in the progeny obtained by crossing w^IR6^ and RevI-H2 flies. Since germline piRNAs are maternally deposited in the embryo and this transgenerational piRNA inheritance triggers piRNA biogenesis in the progeny [14,15], we predicted that if *ZAM*-derived piRNAs in RevI-H2 arose from a *de novo*-formed piRNA cluster, repression operated by this cluster should only be observed when the locus is inherited from the mother. Conversely, if the germline repression of *ZAM* is due to an insertion in a pre-existing piRNA cluster, then piRNA produced by this cluster should induce repression when inherited from either parent (Fig. S4A).

We named ZMD (for maternal deposition of *ZAM*-derived piRNAs) the progeny obtained by crossing a RevI-H2 female and a control male and NZMD (No maternal deposition of *ZAM*-derived piRNAs), the progeny of a RevI-H2 male and a control female. In both crosses, the control line was the line harboring the p*GFP-ZAM* transgene the expression of which is driven in germline cells by the *nanos*-Gal4 driver in the w^1118^ background. In both ZMD and NZMD progenies, the sensor transgene was completely silenced in germline cells, as shown by immunofluorescence and western blot analysis (Fig. 4A-C). This finding suggested that the unknown piRNA cluster that can silence the sensor transgene in the germline does not need maternal deposition of *ZAM*-derived piRNAs to be active. Indeed, the maternal deposition of the general piRNA population, required to activate piRNA clusters in the progeny, was sufficient for the production of *ZAM*-derived piRNAs in the progeny. Therefore, we concluded that the *ZAM*-derived piRNAs produced in the RevI-H2 germline arose from a *ZAM* sequence inserted into a pre-existing germline cluster. To further analyze the sensor silencing and to rule out the possibility that the transgene has become a piRNA cluster by itself, we sequenced and compared ovarian small RNAs from the ZMD progeny (Fig. 4D, right panel) and from a control line in which the p*GFP-ZAM* transgene is expressed in the germline (in the w^IR6^ genetic background: Fig. 4D, left panel and Fig. S4B-C). The results indicated that the sensor transgene was not a *de novo* piRNA cluster because the upstream *GFP* sequence produced very few piRNAs, while a significant amount of piRNAs mapped to the *ZAM* fragment in the ZMD progeny (Fig. 4D; right panel). These data suggested the presence of a new ZAM insertion in a pre-existing germline piRNA cluster.

**Figure 4.**
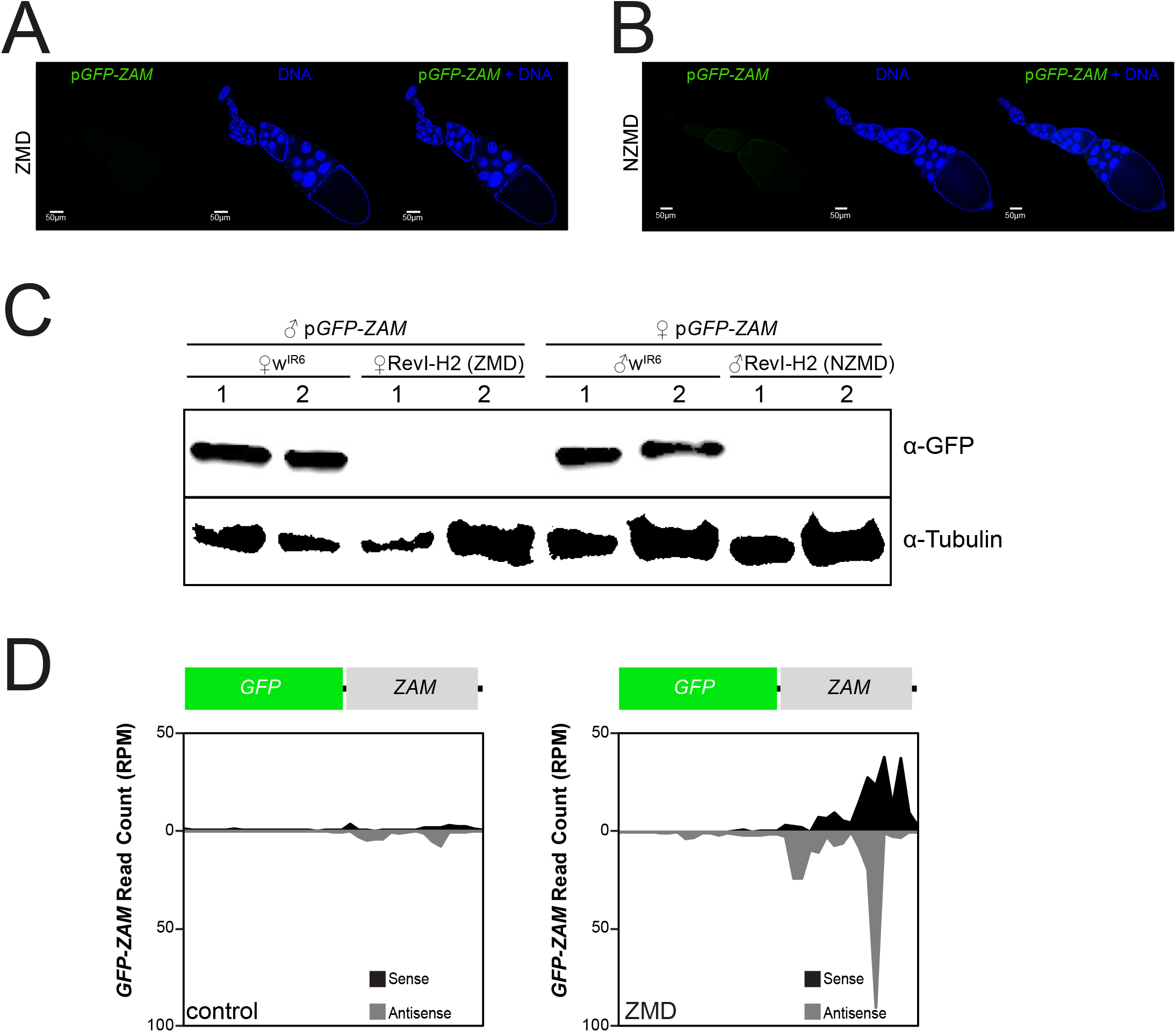
*ZAM*-derived piRNAs are produced from a pre-existing germline piRNA cluster in RevI-H2 ovaries. **a-b.** Confocal images of ovarioles after GFP (green, left panels) and DNA (blue, middle panels) staining. Merged images of the GFP and DNA signals are displayed on the right. Ovarioles were from the progeny of a cross between RevI-H2 females and control males (*ZAM* maternal deposition, ZMD) in *a* and from a cross between RevI-H2 males and control females (No *ZAM* maternal deposition, NZMD) in *b*. In both crosses, the p*GFP-ZAM* line in which ZAM expression is driven in germline cells by a *nanos*-Gal4 driver was the control line. **c** Western blotting of proteins extracted from ovaries of progenies of crosses between w^IR6^ or RevI-H2 and the same control line as in *a* and *b*. The lines used for the crosses are indicated above. Proteins were from two biological replicates (1&2) prepared from 5 pairs of ovaries; α-tubulin was used as loading control. **d** Density profile of piRNAs mapping along the *GFP-ZAM* transgene sequence (allowing up to 3 mismatches). Sense and antisense reads are in black and grey, respectively. The profiles are for crosses between w^IR6^ (left, control) or RevI-H2 (right, ZMD) females and control males harboring the p*GFP-ZAM* transgene.

To genetically map this germline piRNA cluster that produces *ZAM*-derived piRNAs in the germline, we isolated each chromosome of the RevI-H2 line and established three lines harboring (i) the X chromosome from RevI-H2 (X^RevI-H2^; II; III and referred as X^RevI-H2^); (ii) the autosomal chromosome II from RevI-H2 (X; II^RevI-H2^; III and referred as II^RevI-H2^); or (iii) the autosomal chromosome III from RevI-H2 (X; II; III^RevI-H2^ and referred as III^RevI-H2^). It should be noted that the II^RevI-H2^ and III^RevI-H2^ lines carry a wild type *flamenco* locus, while the X^RevI-H2^ line harbors the *flamenco* deletion present in RevI-H2. To identify which chromosome was required for germline production of *ZAM*-derived piRNAs, we assessed the GFP expression of the p*GFP-ZAM* sensor transgene driven by *nanos*-Gal4 in each line. We found that the transgene was silenced in the germline of the X^RevI-H2^ line, like in RevI-H2 (Fig. S4D-E). Conversely, it was expressed in the II^RevI-H2^ and III^Rev I-H2^ germlines (Fig. S4D-E). This indicates that in RevI-H2 ovaries, *ZAM*-derived piRNAs are produced from a germline piRNA cluster localized on the X chromosome.

We sought to identify the precise genomic location of this new *ZAM* insertion in a germline piRNA cluster localized on the X chromosome by performing whole genome sequencing of RevI-H2. We first searched for new *ZAM* insertions in euchromatin within the RevI-H2 genome using the McClintock pipeline [28], which identified seven *ZAM* insertions on the X-chromosome, including the known *ZAM* insertion in the *white* locus at X: 2,799,672..2,799,675. None of these X-chromosome insertions are found in known piRNA clusters. Since the component methods in McClintock do not efficiently detect new TE insertions within repetitive regions, TE nests or piRNA clusters, we used a complementary approach to identify chimeric reads containing both *ZAM* sequence and genomic sequence, which were not uniquely mappable on the reference genome. Using this approach, we identified a new *ZAM* insertion that mapped to a R1 element sequence found at multiple locations within cluster 9 [29], a dual-strand piRNA cluster located next to the X chromosome centromere (Fig. S4F). Analysis of paired-end reads followed by PCR and sequencing in RevI-H2 (Fig. S4G) localized the *ZAM* insertion to one of the three possible sites within this piRNA cluster spanning the interval X: 23,474,449..23,513,109. A previous inverse PCR study in RevI-H2 also detected this *ZAM* insertion but could not map it to a specific genomic location [23]. PCR analysis confirmed that this *ZAM* insertion is absent from the stable lines w^IR6^, ISO1A and w^1118^ (Fig. S4H).

These analyses demonstrated that the RevI-H2 line possesses a *ZAM* insertion in a pre-existing germline piRNA cluster located on the X chromosome.

### Analysis of TEs lost with the *flamenco* deletion in RevI-H2 reveals various patterns of piRNA production

Besides *ZAM*, several other transposons are contained within the *flamenco* deletion in RevI-H2: *Adoxo, Gedeo, Idefix, Phidippo, Pifo, Uxumo* and *Vatovio* (Fig. S1A). To verify whether the genomic deletion also affected the epigenetic regulation of other transposons, we analyzed the piRNA population produced by RevI-H2 ovaries against these different elements. We focused our analysis on *Phidippo* and *Pifo* because they appeared to be mainly silenced by *flamenco*. Indeed, in the control line w^IR6^, *Phidippo* and *Pifo*-derived piRNAs did not harbor a ping-pong signature (Fig. 5A) and were mainly antisense (Fig. 5B-C). Conversely, *Adoxo-, Gedeo-, Idefix*- and *Vatovio*-derived piRNAs displayed a ping-pong signature (Fig. 5A, Fig. S5A). Moreover, 37% of *Phidippo* and 54% of *Pifo*-derived piRNAs that mapped to piRNA clusters [9] mapped *flamenco*, 21% of Phidippo-derived piRNAs mapped cluster 17 (Fig. S5B-C). Notably, cluster 17 has been proposed to be part of the *flamenco* cluster (Zanni et al., 2013), raising the percentage of Phidippo-mapping piRNAs that map to the extended *flamenco* to 58%.

**Figure 5.**
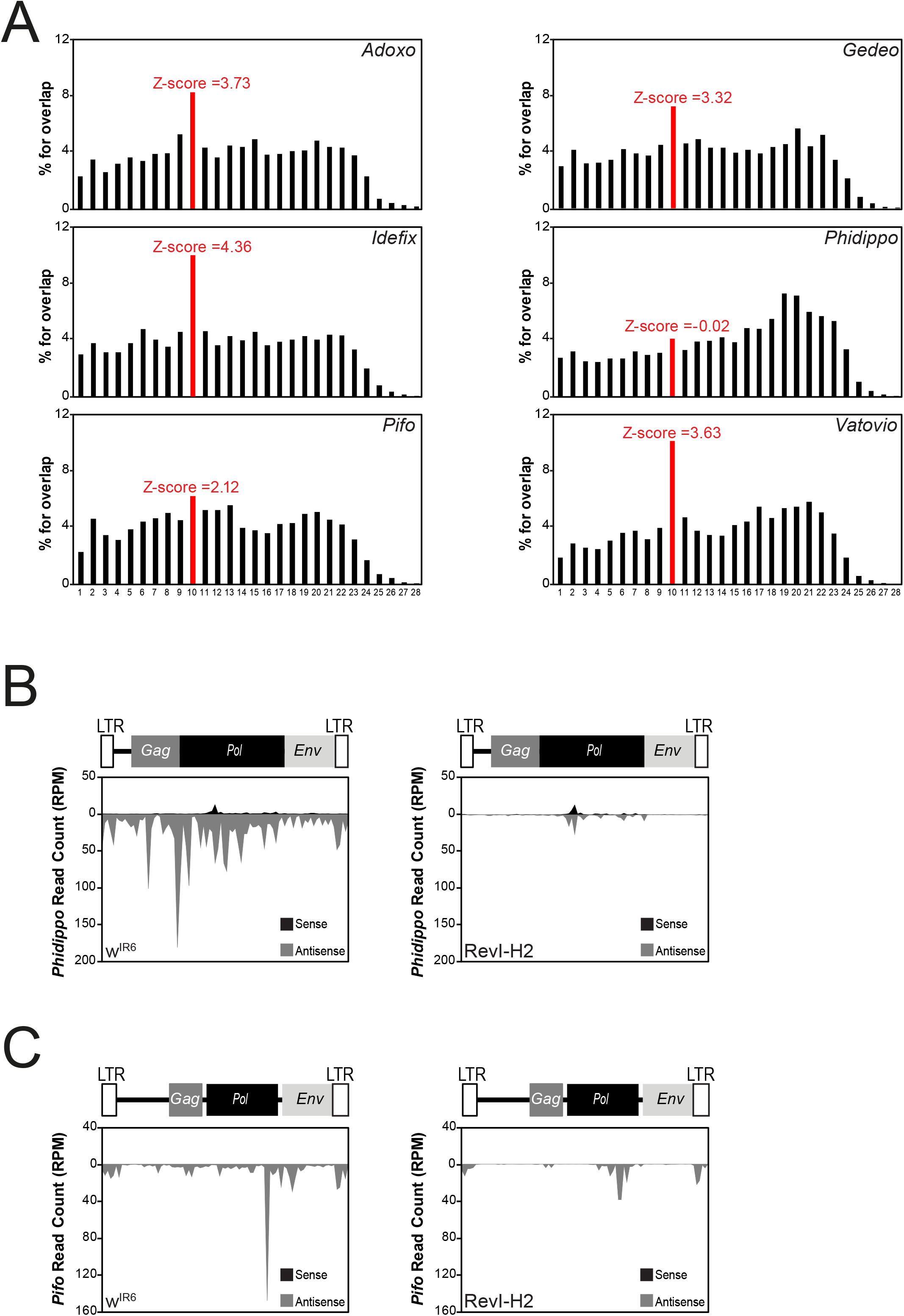
Production of *Phidippo and Pifo*-derived piRNAs is lost in RevI-H2. **a** Histogram for the percentage of 5’-overlaps between sense and antisense *Adoxo-, Gedeo-, Idefix-, Phidippo-, Pifo-* and *Vatovio*-derived piRNAs (23-29nt) in w^IR6^ ovaries. The peak in red defines the 10nt-overlapping pairs and the Z-score is indicated. **b-c** Density profile of *Phidippo-* (*b*) and *Pifo-* (*c*) derived piRNAs along the 7.3 kb *Philippo* sequence and 7.7 kb *Pifo* sequence, respectively, in w^IR6^ (left) and RevI-H2 (right) ovaries (using all piRNAs mapped to the corresponding TE allowing up to 3 mismatches). Sense and antisense reads are represented in black and grey, respectively. The organization of the two TEs is displayed above their respective profile.

In the RevI-H2 line, production of *Phidippo*- and *Pifo*-derived piRNAs was almost abolished (Fig. 5B-C), differently from what observed for *ZAM*-derived piRNAs (Fig. 3D). In contrast to ZAM, which must have an active copy outside the *flamenco* region that gave rise to the new ZAM insertions in RevI-H2 (such as the reference genome copies at 2R:1,808,663..1,817,084 and 3L:24,168,844..24,176,114), no additional active copy of *Phidippo* or *Pifo* has been identified in the reference genome, besides the one in the *flamenco* locus. This indicated that the *Pifo*- and *Phidippo*-derived piRNAs are produced almost exclusively by *flamenco and* that in the absence of additional functional copies, these TEs could not invade the genome, differently from *ZAM*.

### Transposition of *ZAM* in a germline piRNA cluster is an early event

The Rev line was first identified two decades ago [23] based on a phenotypic reversion of the mutated eye phenotype of w^IR6^ flies due to a *de novo ZAM* insertion upstream of the *white* gene. A series of homozygous RevI lines (RevI-H1, RevI-H2 and RevI-H3) were then derived from the initial Rev line. Several secondary mutations affecting eye color were recovered from the initial RevI-H2 line and new lines were successively isolated and called RevII ([30]; see [31] for further description). To further trace when the germline acquired the potential to silence *ZAM*, we sought to determine when the *ZAM* insertion into a germline piRNA cluster occurred. We sequenced ovarian small RNAs from RevII-7 (which was derived 20 years ago from RevI-H2). Detailed analysis of *ZAM*-derived piRNAs in RevII-7 samples showed that *ZAM*-derived sense and antisense piRNAs were produced to an extent similar to what observed in the RevI-H2 line (Fig. 6A). These piRNAs displayed the typical ping-pong signature: a bias for 1U and 10A (Fig. 6B) and the enrichment of 10-nt 5’-overlaps (Fig. 6C). Moreover, 25% of the *ZAM*-derived piRNAs had a PPP with the typical 10A and 1U bias (for the sense and antisense PPPs respectively) (Fig. S6A-F). We concluded that the *ZAM* insertion event into a germline piRNA cluster occurred before the RevII lines were derived from the RevI-H2 line.

**Figure 6.**
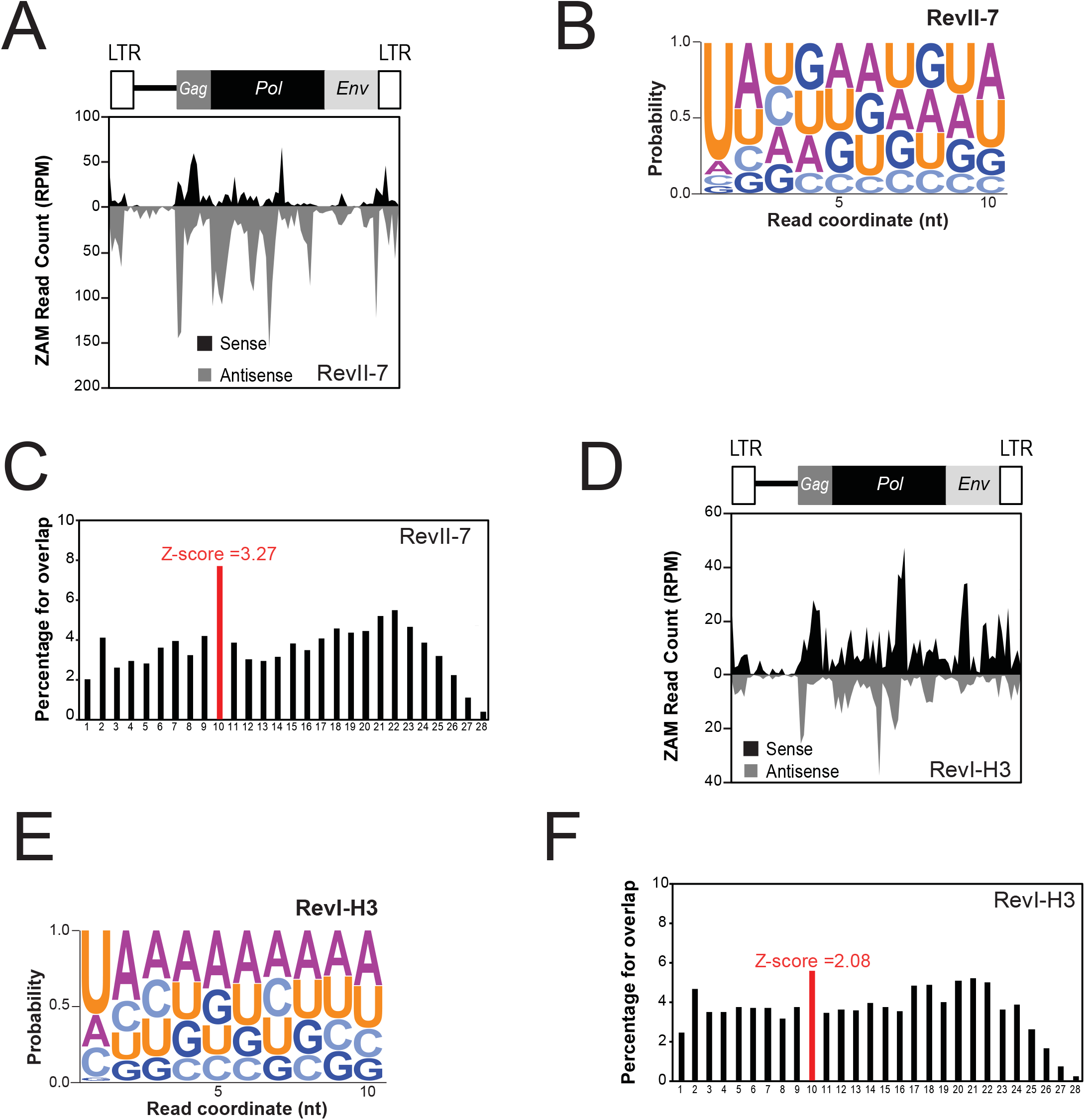
*ZAM* is trapped in a germline piRNA cluster in all analyzed Rev lines. **a** and **d** Density profile of *ZAM*-derived piRNAs along the 8.4Kb *ZAM* sequence in the RevII-7 (*a*) and RevI-H3 (*d*) lines (allowing up to 3 mismatches). Sense and antisense reads are represented in black and grey, respectively. The organization of *ZAM* is displayed above the profiles. **b** and **e** Logo of nucleotide bias for the first ten positions of *ZAM*-derived piRNAs produced in RevII-7 (*b*) and RevI-H3 (*e*) ovaries. The nucleotide height represents its relative frequency at that position. **c** and **f** Histogram showing the percentage of 5’-overlaps between sense and antisense *ZAM*-derived piRNAs (23-29nt) in RevII-7 (*c*) and RevI-H3 (*f*) ovaries. The peak in red defines the 10nt-overlapping pairs and the Z-score is indicated.

Thus, the *ZAM* insertion event may have occurred very early when the three RevI lines (RevI-H1, RevI-H2 and RevI-H3) were established from the initial Rev line. Sequencing of small RNAs from RevI-H3 ovaries and analysis of *ZAM*-derived piRNAs showed again the production of sense and anti-sense piRNAs, but with a high bias for sense piRNAs (Fig. 6D), differently from what observed in the RevII-7 and RevI-H2 lines (Fig. 6A and 3D). The bias for 1U and 10A (Fig. 6E) and the enrichment of the 10-nt 5’-overlap were also present in the RevI-H3 line (Fig. 6F), but to a smaller extent than in the RevI-H2 and RevII-7 lines. In RevI-H3 samples, 20% of the *ZAM*-derived piRNAs possessed a PPP with the typical 10A and 1U bias (Fig. S6B-F). These results suggested that in the RevI-H3 line, which was independently established at the same time as RevI-H2, also carries a *ZAM* insertion in a germline piRNA cluster. However, the differences observed for *ZAM*-derived piRNAs produced in RevI-H2 and RevI-H3 suggested that there are may be secondary changes to the piRNA cluster 9 or that *ZAM* inserted into another piRNA cluster in RevI-H3 different from the one identified in RevI-H2.

In addition to providing context about the timing of the germline invasion, the RevII-7 and RevI-H3 allowed us to determine the conservation of *ZAM* repression over time in independent stocks. To monitor the efficiency of the various *ZAM*-derived piRNAs produced in the germline of the RevII-7 and RevI-H3 lines, we followed the GFP expression of the p*GFP-ZAM* sensor transgene. Like for the RevI-H2 line, the transgene was completely silenced in germline cells and strongly expressed in somatic cells in both RevII-7 and RevI-H3 (Fig. S6G).

To conclude, analysis of the various Rev mutant lines suggested that *ZAM* transposition into a germline piRNA cluster (leading to *de novo ZAM*-derived piRNAs production) is an early and frequent event essential for germline protection against invasion by mobile elements from the surrounding somatic tissue.

## Discussion

TEs have colonized the genome of all living organisms. To ensure their vertical transmission and amplification in multicellular organisms, mobile element transposition has to take place in germ cells. In turn, germ cells have developed specialized strategies to protect the integrity of their genome and thus the species continuity. Using the prototypic somatic element *ZAM* from *D. melanogaster*, we discovered that the germline can rapidly evolve to control the activity of TEs after invasion from the surrounding somatic tissues by trapping copies of the invading element into germline piRNA clusters. This ensures the production of piRNAs against the invading TE and germline genome protection.

### The germline can adapt to the threat of active transposon invasion from surrounding somatic tissues

The *flamenco* locus is a master piRNA cluster, expressed only in somatic follicular cells that do not transfer any genetic information to the progeny. It produces somatic piRNAs characterized by the absence of the ping-pong signature. The very efficient TE silencing in somatic tissue by *flamenco* protects the germline genome against invasion by somatic TEs. The expression pattern of TE-derived piRNAs suggests that several TEs (*gtwin, gypsy, Tabor, gypsy5, gypsy10* and *ZAM*) are almost exclusively controlled by *flamenco*-derived piRNAs [16]. In this study, we demonstrated that in control ovaries, *ZAM* is repressed exclusively in somatic follicular cells and no *ZAM*-derived piRNAs are produced in the germline, leaving the germline genome vulnerable to *ZAM* invasion when its control is lost in somatic follicular cells. In agreement, the p*GFP-ZAM* sensor transgene was not silenced in the germline. This observation also reveals that antisense *ZAM*-derived piRNAs produced in somatic follicular cells are cell autonomous and do not transit to the germline to ensure *ZAM* silencing in this compartment.

In fly ovaries, in addition to the piRNA pathway, the short interfering RNA (siRNA) pathway also is active and involved in TE silencing [32,33]. In addition, it has been reported that, during artificial horizontal transfers of the TE *Penelope* from *D. virilis* to *D. melanogaster*, only 21nt siRNAs are detected in the ovary. However, they cannot completely silence *Penelope* which remained capable of occasional transposition [34]. In the case of *ZAM*, the strong expression of the sensor transgene in the germline cells suggests that neither the siRNA pathway nor any other silencing pathway can silence this TE in the germline.

We previously showed that in the unstable RevI-H2 line in which *ZAM* silencing is released in somatic follicular cells due to a deletion in *flamenco, ZAM* particles produced within follicular cells use the endosomal vitellogenin trafficking system, which is active during late oogenesis, to enter the closely apposed oocyte and invade the germline [22]. At the time of the invasion, no *ZAM*-derived piRNAs were produced in the germline. Therefore, this condition could be compared to what happens when a TE first enters a new species through horizontal transfer [35–38]. For instance, the P element was introduced from *D. willistoni* to *D. melanogaster* by horizontal transfer and a copy of P inserted at the subtelomeric heterochromatin 1A site, which corresponds to a region that gives rise to multiple small RNAs [9,39]. This insertion is sufficient to elicit a strong P repression in *D. melanogaster* P strains [40–42]. Studies on P-M dysgenic hybrid system showed that in the F1 hybrid adult females, the invading paternally inherited P element escapes silencing and mobilizes due to the absence of maternally deposited P-derived piRNAs. With age, fertility is restored and the P element is silenced suggesting also an adaptation to *P* element transposon invasion. However, in contrast to what we observed for *ZAM*, P-derived piRNAs are produced from paternally inherited clusters [43,44]. Our detailed analysis of piRNAs produced by the RevI-H2 ovaries revealed that this line adapted to *ZAM* invasion by trapping a new *ZAM* copy in a germline piRNA cluster, leading to the production of *ZAM*-derived piRNAs in the germline. Hence, the RevI-H2 line is the first example in which the germline, which does not have initially the genetic capacity to produce *ZAM*-derived piRNAs, needs to protect itself from invasion caused by the sudden loss of control of an endogenous somatic cell-specific TE, the expression of which is normally repressed and should not have been a risk for the progeny.

### Retention of invading somatic TEs in germline piRNA clusters protects the germline from further invasion

In this study, we observed the *de novo* production of sense and antisense *ZAM*-derived piRNAs in RevI-H2 ovaries. Analysis of the ZMD and NZMD progenies showed that the piRNA cluster that trapped a *ZAM* copy was activated by maternal deposition of piRNAs other than *ZAM*-derived piRNAs. This finding strongly suggests that the *ZAM* insertion occurred in an existing germline piRNA cluster. The specific features of these *ZAM*-derived piRNAs (10nt overlap and 1U and 10A bias) indicate that they are produced through the germline-specific ping-pong cycle. Moreover, they successfully silenced the p*GFP-ZAM* sensor transgene in germline cells of RevI-H2 ovaries. As *ZAM* is not normally expressed in the germline, the sense transcripts, which are engaged in the ping-pong cycle and produce piRNAs, could arise: (i) from a *ZAM* copy in a germline piRNA cluster, (ii) from dispersed *ZAM* copies inserted in the vicinity of germline promoters, or (iii) from invading *ZAM* mRNAs produced from somatic cells.

Among the 142 piRNA clusters identified in the *D. melanogaster* genome, most of them are significantly enriched in pericentromeric and telomeric heterochromatin [9], regions that concentrate most TEs [45]. We previously proposed a model in which piRNA clusters play the role of TE traps [25]. This model relies on the capacity of TEs to transpose into piRNA clusters that passively acquire new TE content. Thus, TEs that “jump” into piRNA clusters can produce the corresponding piRNAs and silence homologous elements. This mechanism should constitute an adaptive advantage that can then be fixed by evolutionary selection. How piRNA clusters are formed and then produce piRNAs to repress a novel invasive TE is not well understood yet. Our findings indicate that *de novo* piRNAs can be produced by germline cells after *ZAM* invasion from another cellular lineage (*i.e*. somatic follicular cells) and successfully counteract the invasion. This suggests that invasive TEs can be trapped by piRNA clusters. *ZAM* trapping into a pre-existing piRNA cluster could result from a random transposition event. However, we found that in all the Rev lines analyzed, a germline piRNA cluster trapped a *ZAM* copy. Therefore, TE trapping by piRNA clusters seems to be a frequent event and/or there is selective pressure to maintain a newly inserted *ZAM* copy in a germline piRNA cluster. The chromatin structure or some physical constraints, such as the nuclear organization of piRNA clusters in the genome, may play a role in transposon trapping. It has been suggested that in *Arabidopsis thaliana*, a nuclear structure, termed KNOT, in which TE-enriched regions of all five chromosomes are entangled, is a preferential insertion site for TEs [46]. In addition, the low recombination rate of these heterochromatic regions might facilitate TE accumulation for further development into piRNA clusters [47].

## Conclusion

In our model system, *ZAM* internal invasion of the germline from another cell type mimics a TE horizontal transfer. This constitutes a unique opportunity to investigate the germline behavior after TE invasion in a system that experimentally imitates evolution. However, we cannot exclude that *ZAM* silencing is progressive, thus requiring several generations for complete repression. Finally, it is thought that piRNA clusters allow germ cells to record the TEs to which they have been exposed to over time, resulting in their silencing by the piRNA pathway. For this reason, the content of all piRNA clusters could be considered as the genetic vaccination record of that fly line or population.

## Methods

### Fly stocks, transgenic lines and crosses

All experiments were performed at 20 °C. The strains *nanos*-Gal4, *actin*-Gal4, w^1118^, w^IR6^ and the various Rev lines [24,48] were from the GReD collection. The FM7c (#2177) strain, the RNAi lines against *white* (#35573), *Aub* (#35201) and *AGO3* (#35232) were from the Bloomington Drosophila Stock Center. The p*GFP-ZAM* sensor transgene (located on chromosome 2) was generated by inserting part of the *ZAM env* region into the UASp-*GFP* vector containing F_LP1_ Recombination Target (FRT) sequences [49] after *Not*I/*Bam*HI digestion. The *ZAM env* region was amplified by Taq polymerase using the primers 5’-GAAGCGGCCGCCGGGACTCACGACTGATGTG-3’ and 5’-GAAGGATCCCGGAGGAATTGGTGGAGCGA-3’. The FRT-ZAM-FRT construct is in sense orientation relative to the *GFP* gene. Gal4-driven p*GFP-ZAM* sensor lines were established by crossing the p*GFP-ZAM* line with the *actin*-Gal4 or *nanos*-Gal4 driver lines.

### Immunofluorescence

Ovaries from 3-to 5-day-old flies were dissected in Schneider’s Drosophila Medium, fixed in 4% formaldehyde/PBT for 15min, rinsed three times with PBT (1X PBS, 0.1% Triton, 1% BSA), incubated in PBT for at least 1h and then with goat anti-GFP (ab5450, Abcam; 1/1000), mouse anti-Ago3 (1/500) [9] or rabbit anti-Aub (1/500) [9] antibodies overnight. After 3 washes in PBT, ovaries were incubated with the corresponding secondary antibodies (1/1,1000), coupled to Alexa-488, Cy3 or Alexa-488 respectively, for 90min. After two washes, DNA was stained with the TOPRO-3 stain (1/1,000). Three-dimensional images were acquired on Leica SP5 and Leica SP8 confocal microscopes using a 20X objective and analyzed using the Fiji software [50]. Images of the progeny of w^IR6^ and Rev crosses were processed with the same parameters.

### Protein extraction and western blotting

At least 5 pairs of ovaries from 3-to 5-day-old flies were dissected in 200μl of Lysis Buffer (17.5mM HEPES, 1.3mM MgCl_2_, 0.38M NaCl, 0.18mM EDTA, 22% glycerol, 0.2% Tween-20 and protease inhibitor cocktail from Roche). After sonication, supernatants were recovered and 400μg of proteins were loaded on precast 4-15% acrylamide gels. Western blots were probed using anti-GFP (Ozyme; #JL-8; 1/1,000) and anti-tubulin (to confirm equal loading) (Sigma, #DM1A, 1/5,000) antibodies, followed by an anti-mouse (Abliance; 1/1,000) secondary antibody and then the Clarity Western ECL reagent (BioRad). Densitometric analysis was performed on non-saturated signals using the Image Lab™ software (BioRad).

### Small RNA sequencing and bioinformatics analysis of piRNAs

Total RNA was isolated from 80-100 pairs of ovaries from 3-to 5-day-old flies or from ovarian somatic sheath (OSS) cell culture (for analysis of piRNA production by somatic follicular cells) with TRIzol Reagent (Ambion). After 2S RNA depletion, deep sequencing of 18-30nt small RNAs was performed by Fasteris S.A. (Geneva/CH) on an Illumina Hi-Seq 4000. Illumina small RNA-Seq reads were loaded into the small RNA pipeline sRNAPipe [51] for mapping to the various genomic sequence categories of the *D. melanogaster* genome (release 6.03). All libraries were normalized to the total number of genome-mapped reads (no mismatch). For the analysis, 23-29nt RNAs were selected as piRNAs. All the analyses were performed using piRNAs mapped to TEs (0 to 3 mismatches) or genome-unique piRNAs mapped to piRNA clusters, as defined by [9] (no mismatch allowed), the strand relative to the transposon or the genome being determined [9]. The window size was of 428nt for *flamenco*, 91nt for *ZAM*, 80nt for *Burdock*, 87nt for *Pifo and* 85nt for *Phidippo* to establish the density profile of piRNAs and depended of the TE size. The ping-pong signature was assessed by counting the proportion of sense piRNAs with an overlap of 10nt with antisense piRNAs, based on piRNAs mapping to the analyzed TE (0 to 3 mismatches). The proportions of 1 to 28nt-long overlaps were determined and the percentage of 10nt overlaps defined as ping-pong signature. The Z-score was determined on the proportions of 1 to 23nt-long overlaps and considered significant for values >1.96. The nucleotide frequency for each position within the 10nt-overlap was determined for the piRNAs mapping to the analyzed TE (0 to 3 mismatches) with ping-pong partners. Logos were generated with the WebLogo web server [52].

### Genome sequencing and analyses of new ZAM insertions in RevI-H2

Genomic DNA from RevI-H2 was extracted from a sample of mixed sex adult flies using standard protocol. Input DNA was tagmented using the Illumina Nextera DNA sample preparation kit (Cat. No. FC-121-1030). Following a cleanup using the Zymo-Spin kit (Cat. No. D4023), the purified, tagmented DNA was then amplified *via* limited-cycle PCR that also added the indices (i7 and i5) and sequencing primers. AMPure XP beads (Cat. No. A63881) were then used to purify and size select the library DNAs. The libraries were then normalized to 2nM and pooled prior to cluster generation using a cBot instrument. The loaded flow-cell was then paired-end sequenced (2×101 bp) on an Illumina HiSeq2500 instrument.

To identify new ZAM insertions in the RevI-H2 genome, we used two complementary approaches. First, we used the McClintock system which aims to identify non-reference TE insertions using multiple component TE detection systems (commit: 9f53a5b4e1fc977b22a77babfb24461face407d3, options -m “popoolationte retroseq temp ngs_te_mapper te-locate”). Because McClintock component methods do not efficiently detect new TE insertions within repetitive regions, we developed a second approach to identify candidate TE insertions in piRNA clusters. Chimeric reads containing genomic sequence and *ZAM* 5’- or 3’-sequence were isolated from the unmappable reads. *ZAM* sequences were then stripped off from these chimeric reads and the resulting flanking sequences mapped to the *D. melanogaster* Release 6.03 genome. This approach identified a novel *ZAM* insertion in piRNA cluster 9 as defined by [29]. We validated the presence of the insertion by PCR on DNA extracted from RevI-H2 flies. The following primers were used for Fig. S4G: primer F 5’-CTCACCATTTCCTCCTTGAC-3’ and primer R 5’-CTCCCAATCATCTCCTCCAA-3’. Sequencing of the amplicon was done by GATC Biotech.

## Declarations

### Acknowledgments

We thank Nathalie Guéguen, Françoise Pellissier and Nadège Anglaret for technical assistance. We thank Arpita Sarkar who constructed the plasmid *pGFP-ZAM*, Marion Delattre for assistance sequencing the w^IR6^ and RevI-H2 small RNA libraries, J. Brennecke for Aub and AGO3 antibodies.

We thank all members of the team for discussion and critical comments.

### Availability of data and material

RNA-seq and DNA-seq datasets supporting the conclusions of this article are available in the NCBI database, PRJNA483852 (https://www.ncbi.nlm.nih.gov/sra/SRP155919).

### Authors’ contribution

EB, CD conceived the study. EB, CD, MY, NM designed and performed experiments. EB, CD gathered and analyzed small RNA-seq data. CD, EB and SJ performed bioinformatic analyses. CB and SJ gathered and analyzed the DNA-seq data. EB and CD analyzed the data. SJ and CV participated in discussions about the project and critically read the manuscript. CD and EB wrote the article. All authors read and approved the final manuscript.

### Ethics approval and consent to participate

Not applicable.

### Consent for publication

Not applicable.

## Supplemental Figure Legends

**Figure S1. The RevI-H2 line carries a deletion that removes *ZAM* from the *flamenco* piRNA cluster. a** Structure of the *flamenco* piRNA cluster in *D. melanogaster*. TEs located in the region of *flamenco* deleted in RevI-H2 are presented individually above *flamenco* according to [25]. The centromere of the X chromosome is on the right-hand (proximal) side. Sense-strand transcription for *flamenco* and TE orientation are indicated by black arrows. The *flamenco* deletion distal breakpoint in RevI-H2 [25] is indicated by a red arrow. The chromosome coordinates are according to release 5 of the *D. melanogaster* genome. **b** Genome browser panel showing *flamenco* piRNA levels in w^IR6^ and RevI-H2 line. The refined Release 6.03 coordinate for the breakpoint of the RevI-H2 deletion in *flamenco* reported in [25] is displayed by a red line.

**Figure S2. In RevI-H2 ovaries, piRNAs derived from *Burdock*, the prototypic germinal TE, present similar features as those derived from *ZAM*. a-b** Logo of nucleotide bias for the first ten positions of sense (*a*) and antisense (*b*) *ZAM*-derived piRNAs with ping-pong partner (PPP) produced in RevI-H2 ovaries. The nucleotide height represents its relative frequency at that position. **c** Density profile of *Burdock*-derived piRNAs along the 6.4kb *Burdock* sequence in w^IR6^ (left) and RevI-H2 (right) ovaries (all mapping piRNAs allowing up to 3 mismatches). Sense and antisense reads are in black and grey, respectively. The organization of *Burdock* is displayed above the profiles. **d** The total amount of *Burdock*-derived piRNAs produced in w^IR6^ and RevI-H2 ovaries was quantified from the profiles in *c*. **e** Histogram showing the percentage of 5’-overlaps between sense and antisense *Burdock*-derived piRNAs (23-29nt) in w^IR6^ (top) and RevI-H2 (bottom) ovaries. The peak in red defines the proportion of 10nt-overlapping pairs and the Z-score is indicated. **f** Bar diagram indicating the percentage of *Burdock*-derived piRNAs with ping-pong partner (PPP) in the w^IR6^ and RevI-H2 lines. **g** Analysis of the nucleotide bias for sense (+) and antisense (−) *Burdock*-derived piRNAs with PPP in w^IR6^ (left) and RevI-H2 (right). The percentages of PPPs with a 1U and 10A are displayed. **h** Analysis of nucleotide bias for sense (+) and antisense (−) *ZAM*- and Burdock-derived piRNAs with PPPs in RevI-H2 ovaries.

**Figure S3. *ZAM*-derived piRNAs are *de novo* produced by the germline of RevI-H2 ovaries. a-b** Confocal images of ovarioles from w^IR6^ (*a*) and RevI-H2 (*b*) ovaries after labeling with anti-Aub (green, top) and -Ago3 (red, middle) antibodies and DNA (blue, bottom) staining. Merged images of the Aub or Ago3 signal and DNA staining are displayed on the right panels. **c** Density profile of *ZAM*-derived piRNAs along the *ZAM* sequence produced in early embryos from RevI-H2 (allowing up to 3 mismatches). Sense and antisense reads are represented in black and grey, respectively. *ZAM* organization is displayed above the profile. **d** The total amount of *ZAM*-derived piRNAs produced in early embryos from RevI-H2 was quantified from the profile in Fig. S3C. Sense and antisense reads are represented in black and grey, respectively. **e** Histogram showing the percentage of 5’-overlaps between sense and antisense *ZAM*-derived piRNAs (23-29nt) in early embryos from RevI-H2. The peak in red defines the proportion of 10nt-overlapping pairs and the Z-score is indicated. **f** Bar diagram indicating the percentage of *ZAM*-derived piRNAs with ping-pong partner (PPP) in early RevI-H2 embryos. **g** Analysis of the nucleotide bias for sense (+) and antisense (−) *ZAM*-derived piRNAs with PPPs in early RevI-H2 embryos. The percentages of PPPs with a 1U and those with a 10A are displayed. **h** Confocal images of ovarioles after GFP (green, left panels) and DNA (blue, middle panels) staining. Ovarioles were from the progeny of a cross between w^IR6^ or RevI-H2 females and males carrying the p*GFP-ZAM* sensor transgene – obtained after excision of the *ZAM* sequence upon recombination between the flanking FRTs – driven by *actin*-Gal4. Right panels, merged images of GFP and DNA labeling. **i-j** Confocal images of ovarioles after GFP (green), Aub (red, *i*) or AGO3 (red, *j*) and DNA (blue) staining. Ovarioles were from the progeny of a cross between RevI-H2 females carrying the *nanos*-Gal4 driver and males carrying the p*GFP* sensor transgene with either of the RNAi *Aub*- or *AGO3*-KD (Knock-Down). The *white*-KD was used as a control. Right panels, merged images of GFP, Aub (*i*) or AGO3 (*j*) and DNA labeling.

**Figure S4. *ZAM*-derived piRNAs originate from a germline piRNA cluster localized on the X chromosome. a** Schema representing crosses performed to generate the ZMD and NZMD F1 progeny analyzed in Figure 4. The schema details both hypothesis for the new *ZAM* insertion in the RevI-H2 line. Only the X chromosome is displayed on the schema. The control line is the p*GFP-ZAM* line in which expression is driven in germline cells by a *nanos*-Gal4 driver. **b-c** Confocal images of ovarioles after staining for GFP (green, left panels) and DNA (blue, middle panels). Merged images of the GFP and DNA signals are on the right. Ovarioles were from the progeny of a cross between w^IR6^ females and control males (*b*) and from the reciprocal cross between w^IR6^ males and control females (*c*). In both crosses, the p*GFP-ZAM* line in which expression is driven in germline cells by a *nanos*-Gal4 driver was the control line. d Western blot analysis of proteins extracted from ovaries of progenies of crosses between X^Rev^, II^Rev^ or III^Rev^ females with control males. The control line was the same as in *b* and *c*. Proteins were from two biological replicates (1&2) prepared from 5 pairs of ovaries and α-tubulin was the loading control. **e** Confocal images of ovarioles after staining for GFP (green, left panels) and DNA (blue, middle panels). Merged images of the GFP and DNA signals are on the right. Ovarioles were from the progeny of a cross between X^Rev^, II^Rev^ or III^Rev^ females with control males. The control line was the same as in *b* and *c*. **f** Structure of the region where the new *ZAM* insertion was identified in RevI-H2. The insertion is located in the dual-strand piRNA cluster 9 (according to ranking of piRNA clusters identified in *D. melanogaster* by [29]). *ZAM* is in genomic minus strand orientation. The chromosome coordinates are according to release 6 of the *D. melanogaster* genome. The *su*(*f*) genetic marker is indicated. The pericentromeric part of the X chromosome is displayed by a black box, euchromatin as a black line, *flam* and the other piRNA clusters by colored boxes. Primers (F and R) used to confirm the presence of the *ZAM* insertion by PCR (Fig. S4H) are depicted by red arrows. The *ZAM* insertion was localized in region X: 23,474,449..23,513,109 by genome sequencing of RevI-H2 (3 identical possible insertion sites). **g** The sequenced junctions between *ZAM* and the genomic flanks. The *ZAM* sequence is displayed in black capital letter and the genomic flanking sequence matching to cluster 9 in orange lower case letters. Target site duplications (TSD) are boxed. **h** PCR analysis of the new *ZAM* insertion identified in RevI-H2. Used primers are displayed in Fig. S4F. Only RevI-H2 presents an amplicon showing that the w^IR6^, Iso1A and w^1118^ lines are devoid of this insertion and confirming the presence of this *ZAM* insertion located in cluster 9 in RevI-H2.

**Figure S5. *Phidippo*- and *Pifo*-derived piRNAs are mainly produced by the *flamenco* cluster. a** Logo of nucleotide bias for the first ten positions of *Adoxo-, Gedeo-, Idefix*- and *Vatovio*-derived piRNAs with ping-pong partner (PPP) produced in w^IR6^ ovaries. The nucleotide height represents its relative frequency at that position. **b** and **c** Pie charts showing the proportion of *Phidippo-* (*a*) and *Pifo*-derived piRNAs (*b*) mapped (allowing up to 3 mismatches) to the 142 piRNA clusters (allowing no mismatch, piRNA clusters as in [9]) in the w^IR6^ line. **d** Pie charts showing the proportion of *Adoxo-, Gedeo-, Idefix-* and *Vatovio*-derived piRNAs mapped (allowing up to 3 mismatches) to the 142 piRNA clusters (allowing no mismatch, piRNA clusters as in [9]) in the w^IR6^ line.

**Figure S6. *ZAM*-derived piRNAs produced in the different Rev lines display similar features. a** Percentage of *ZAM*-derived piRNAs with ping-pong partner (PPP) in the RevII-7 and RevI-H3 lines. **b** Analysis of the nucleotide bias for sense (+) and antisense (−) *ZAM*-derived piRNAs with PPPs in the RevII-7 and RevI-H3 lines. The percentages of PPPs with a 1U and a 10A are shown. Both lines had a 10A bias for sense piRNAs and a 1U bias for antisense piRNAs. **c-f** Logo of nucleotide bias for the first ten positions of sense (*c,e*) and antisense (*d, f*) *ZAM*-derived piRNAs with ping-pong partner (PPP) produced in RevI-H3 (*c-d*) and RevII-7 (*e-f*) ovaries. The nucleotide height represents its relative frequency at that position. **g** Confocal images of ovarioles after staining for GFP (green, left panels) and DNA (blue, middle panels). Merged images of the GFP and DNA signals are on the right. Ovarioles were from the progeny of crosses between w^IR6^ (top), RevII-7 (middle) or RevI-H3 (bottom) females with males that harbor the p*GFP-ZAM* sensor transgene driven by the *actin*-Gal4 driver. **h** Western blot analysis of proteins extracted from ovaries of progenies of crosses between w^IR6^, RevI-H2, RevII-7 or RevI-H3 females with a control male. The p*GFP-ZAM* line in which *ZAM* expression is driven in germline cells by a *nanos*-Gal4 driver was the control line. Proteins were from two biological replicates (1&2) prepared from 5 pairs of ovaries and α-tubulin was the loading control.

