## Supplementary figures and images for "Trapping a somatic endogenous retrovirus into a germline piRNA cluster immunizes the germline against further invasion"

### FigureS1

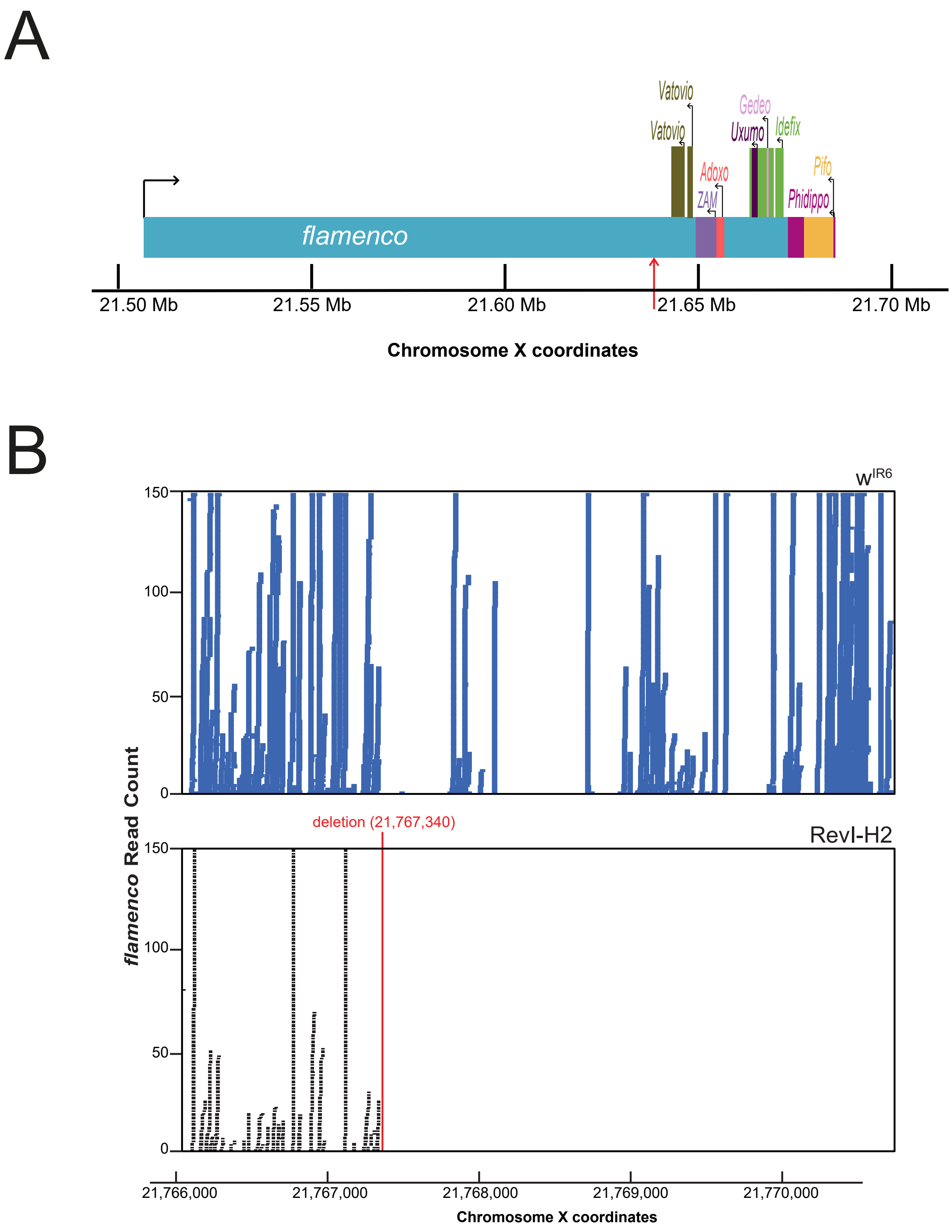

### FigureS2

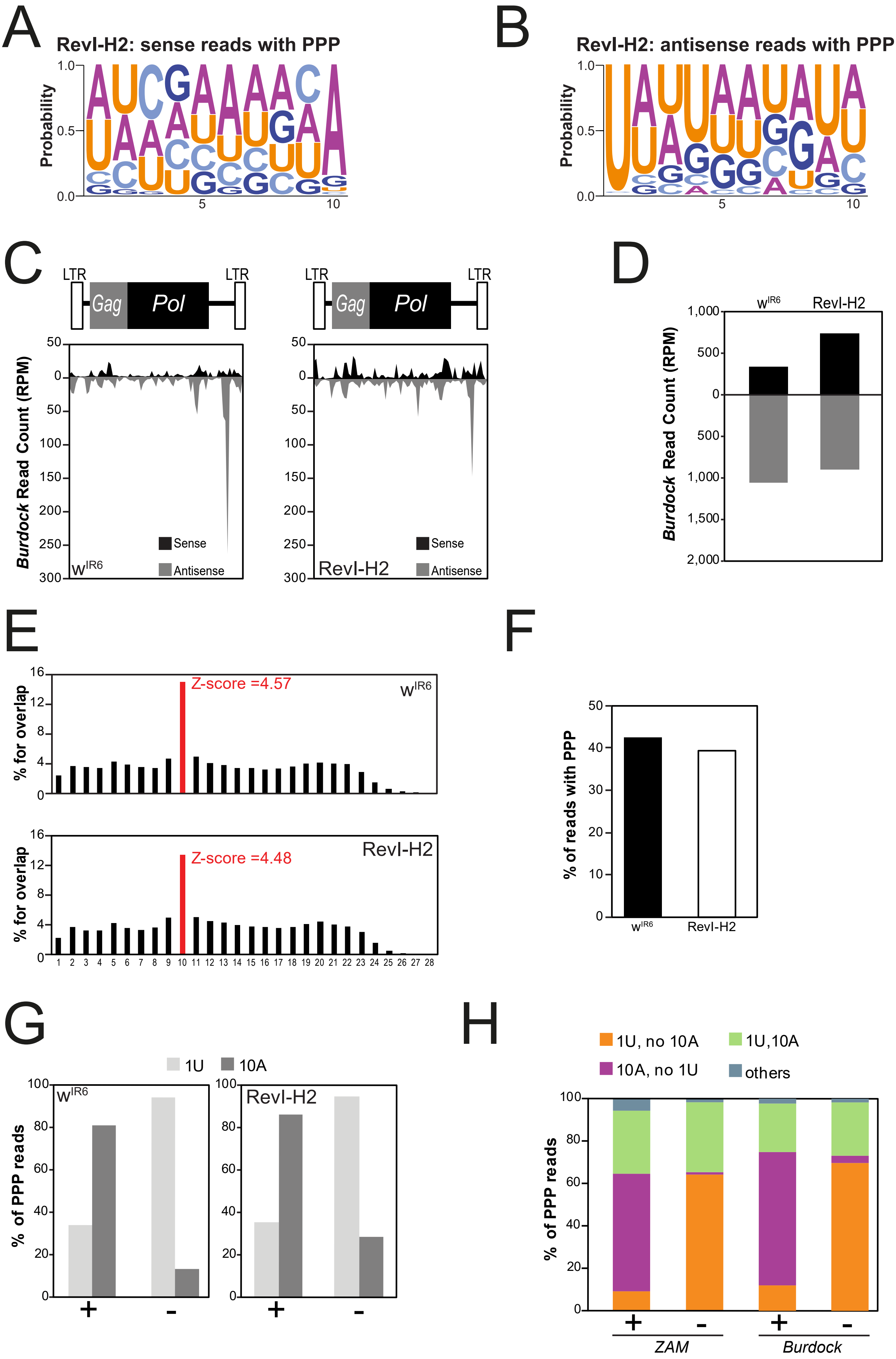

### FigureS3

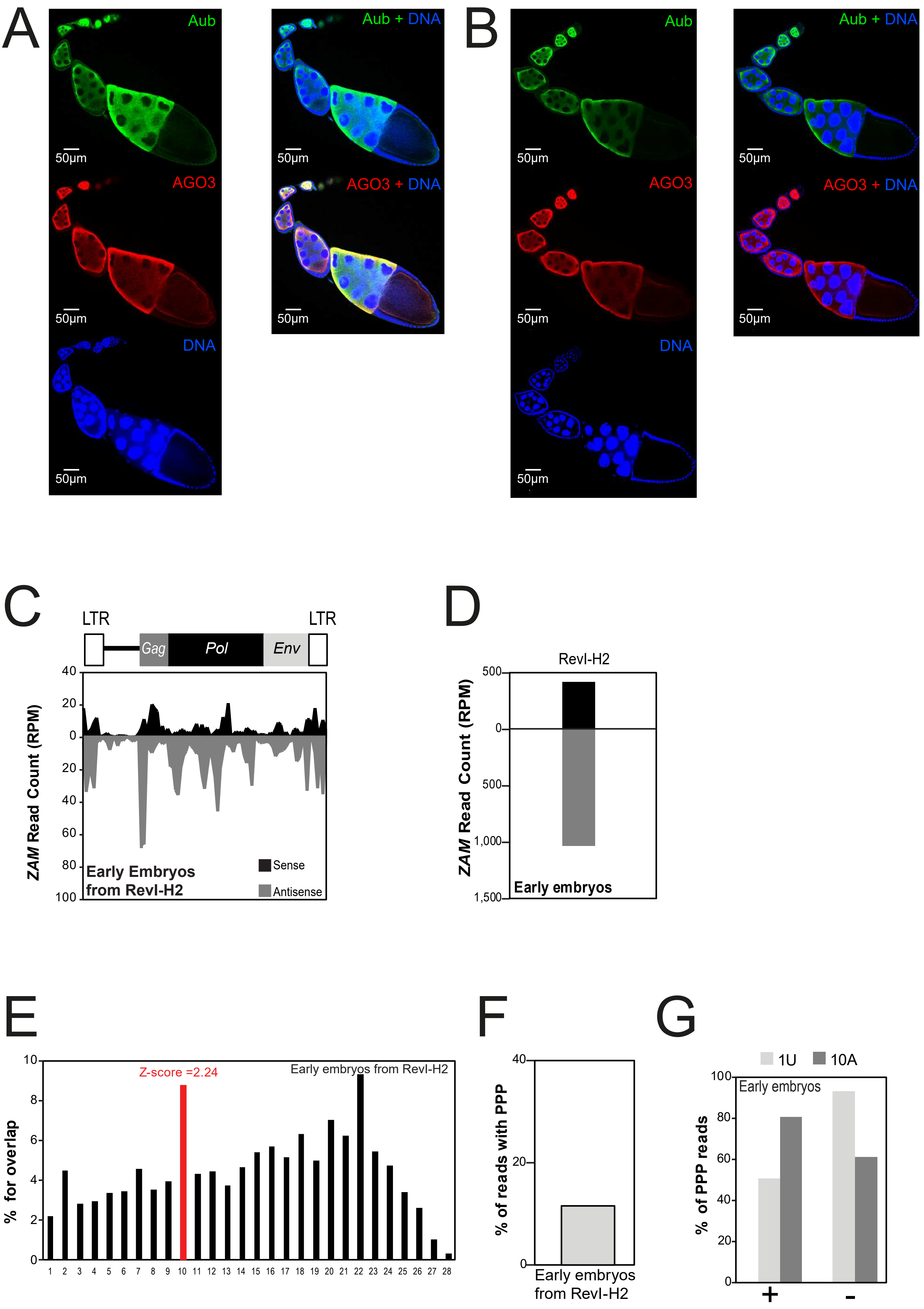

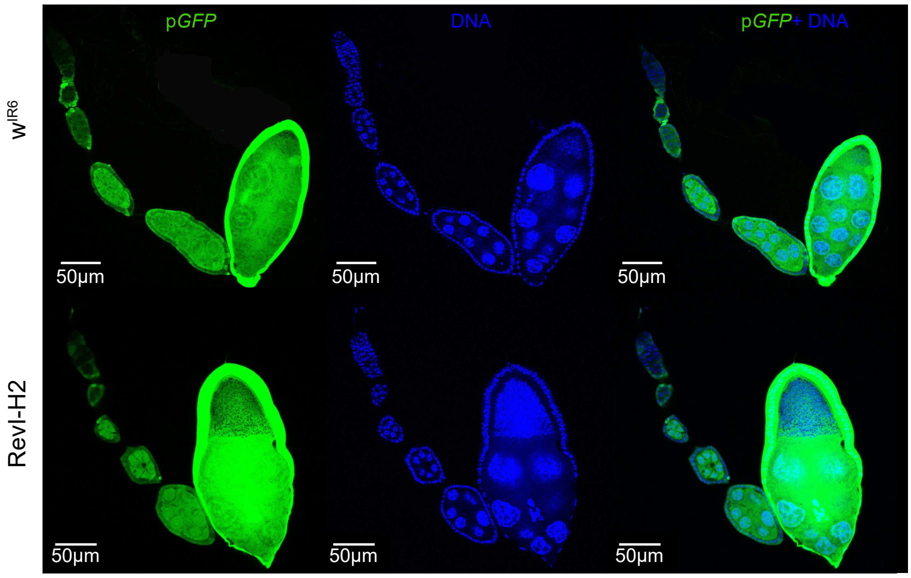

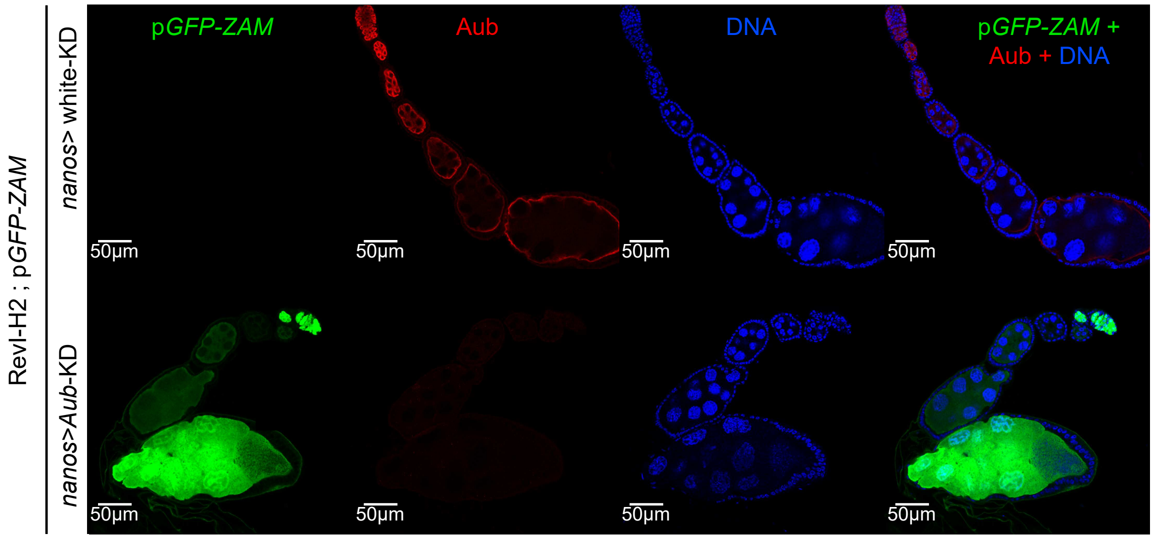

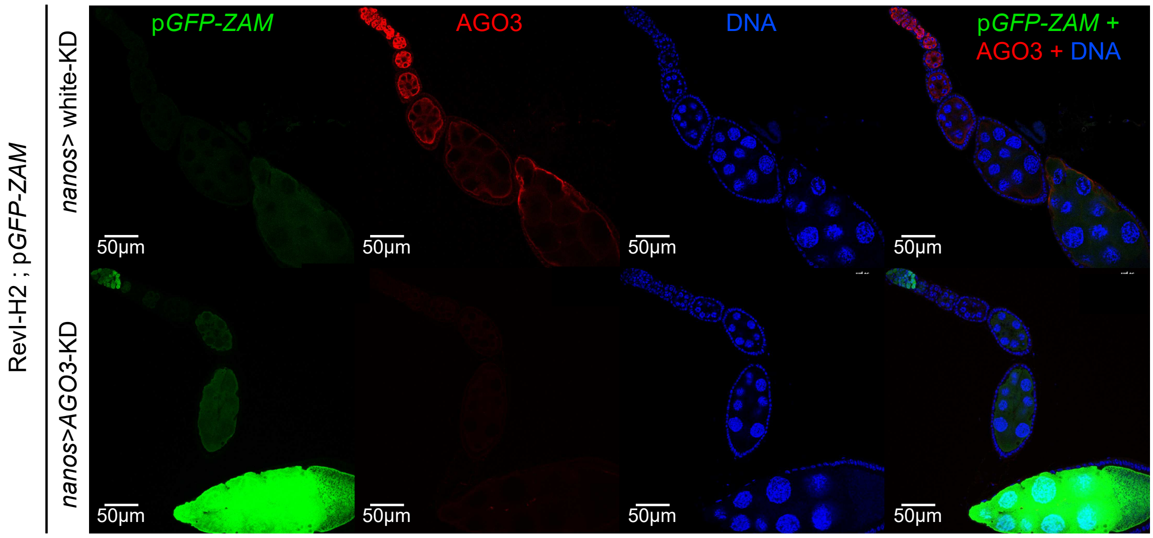

### FigureS4

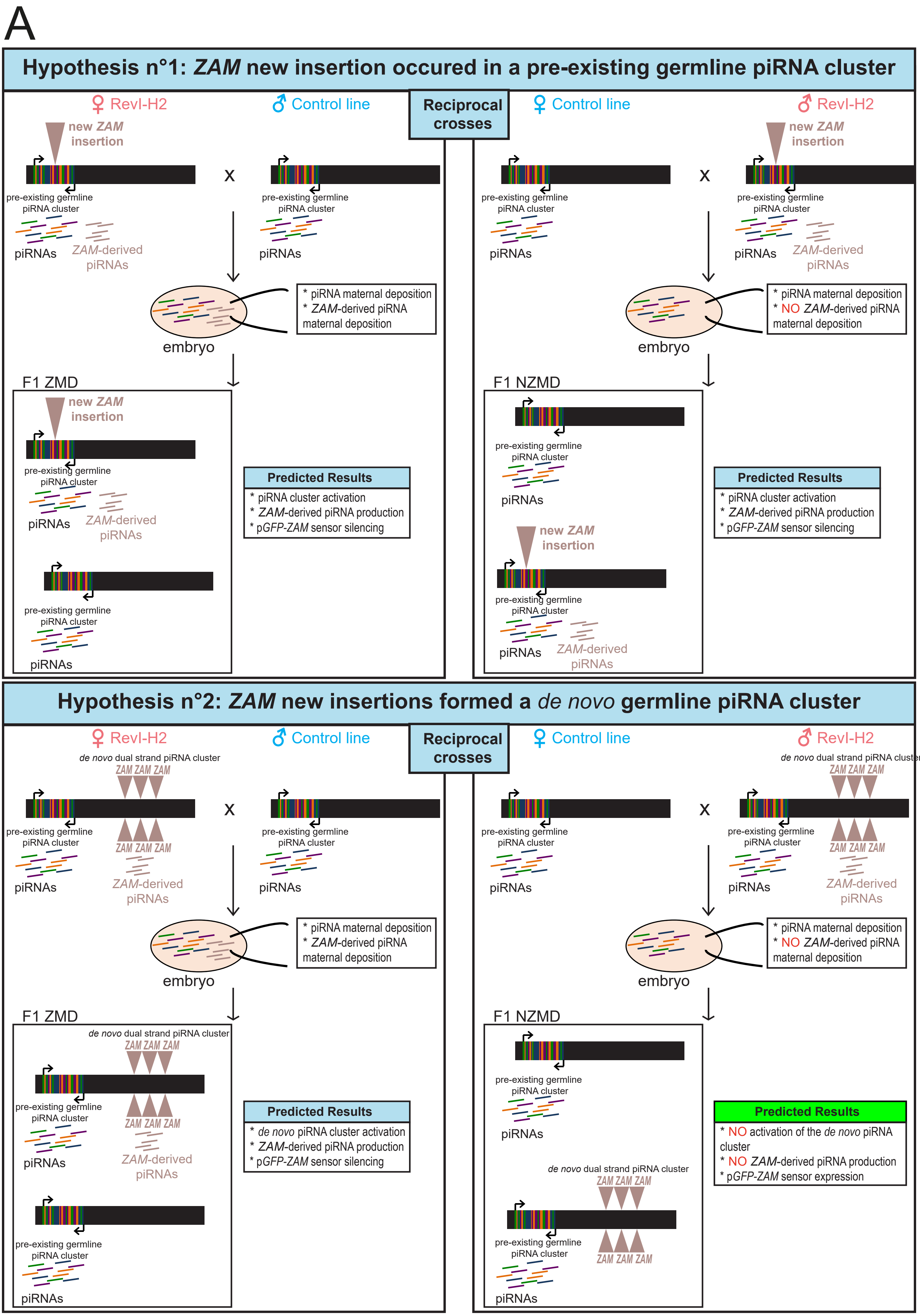

B

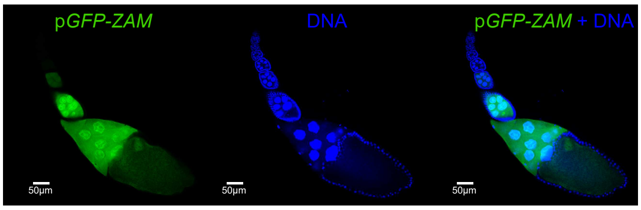


F1 progeny (cross female w$^{IR6}$ with male p*GFP-ZAM*)

C

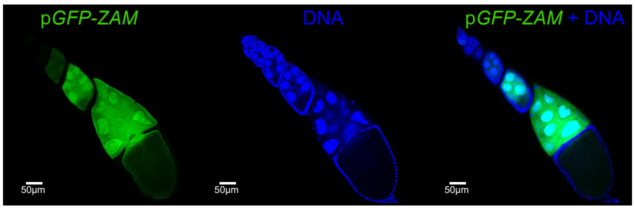


F1 progeny (cross male w$^{IR6}$ with female p*GFP-ZAM*)

D

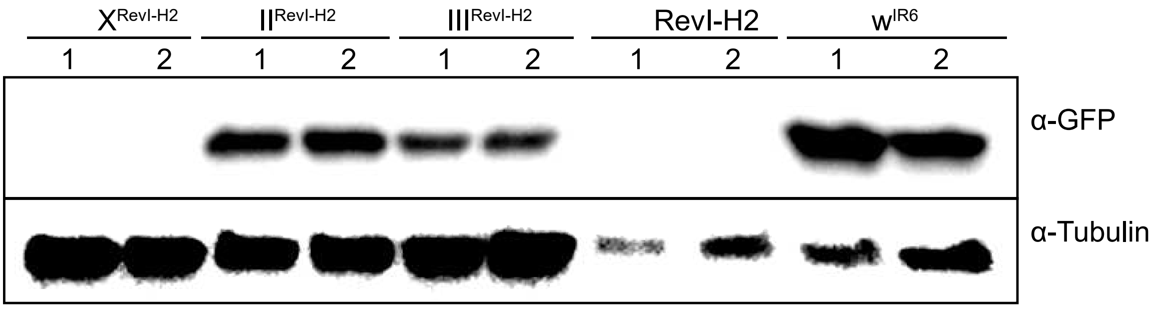


E

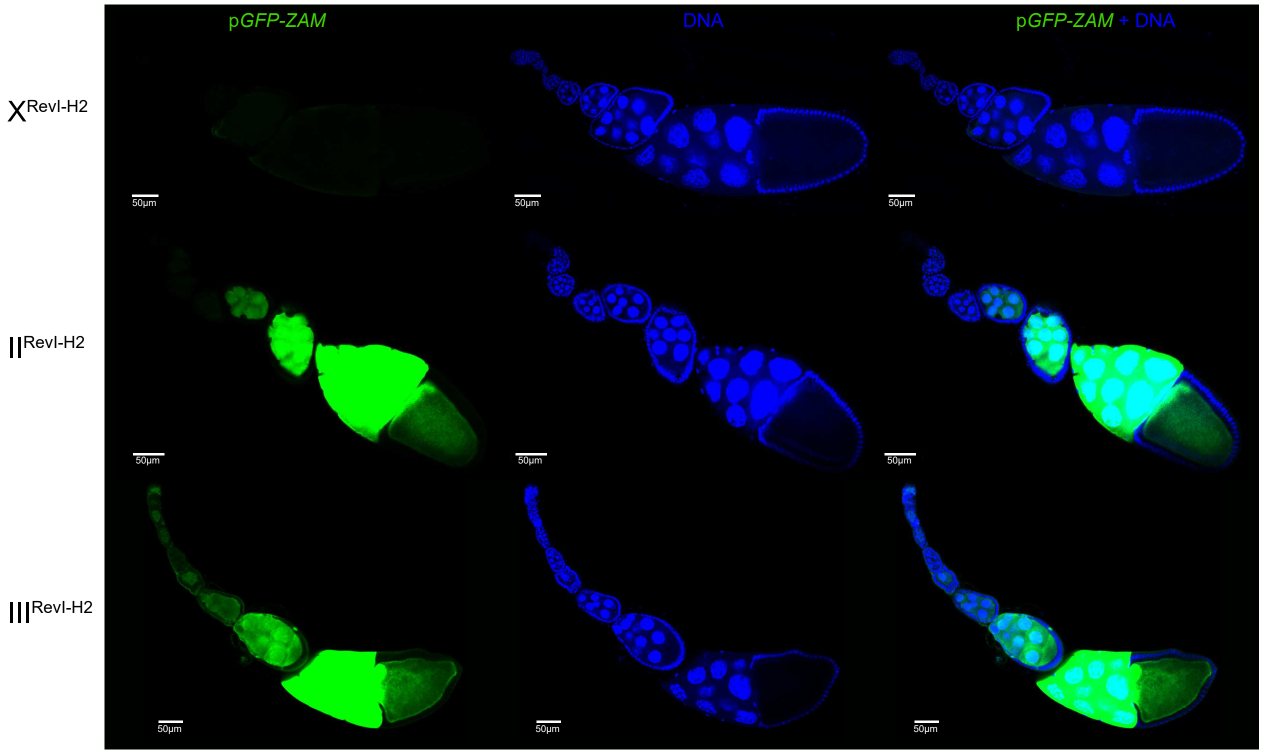

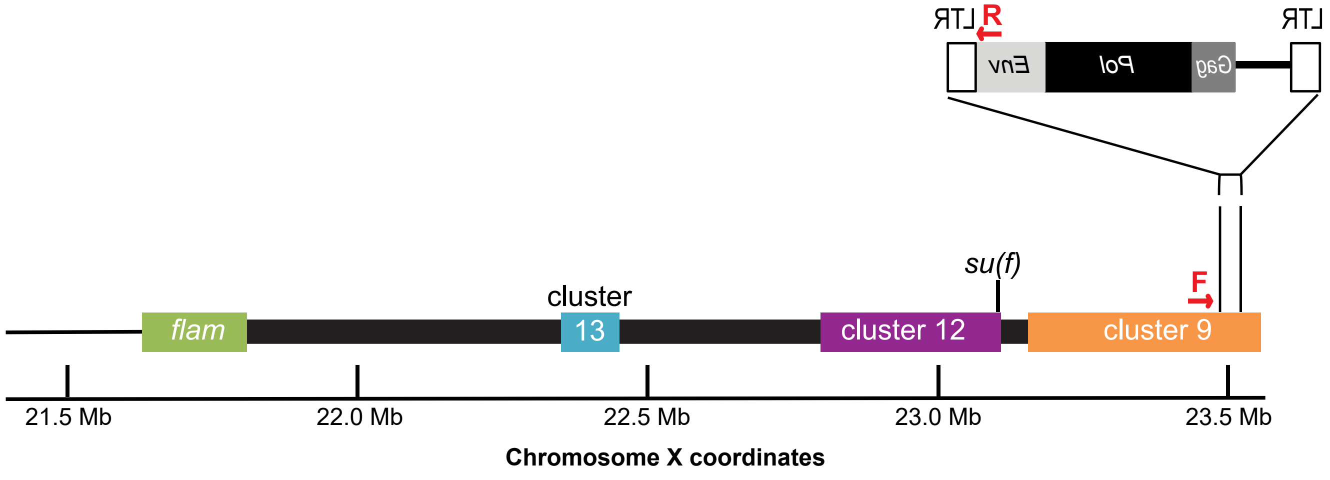

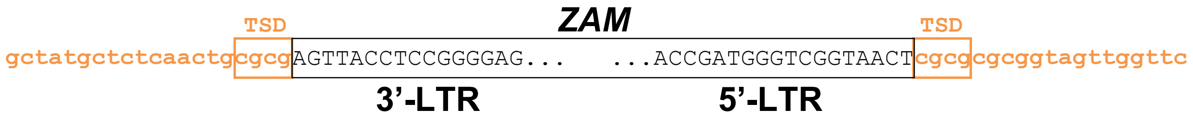

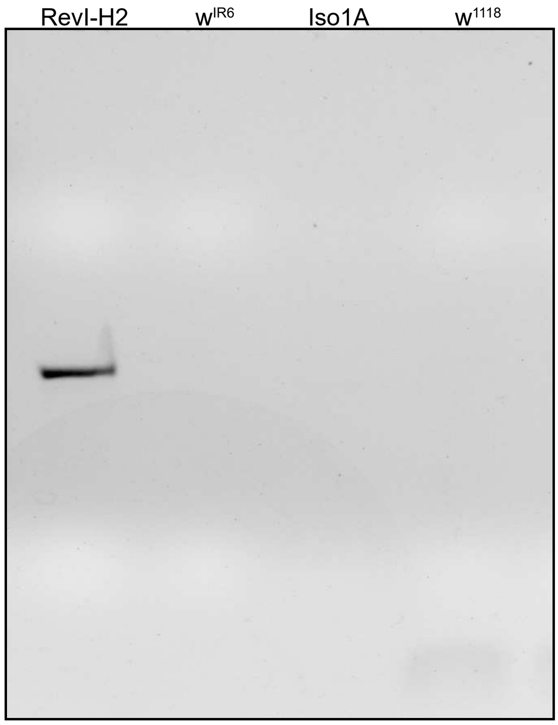

### FigureS5

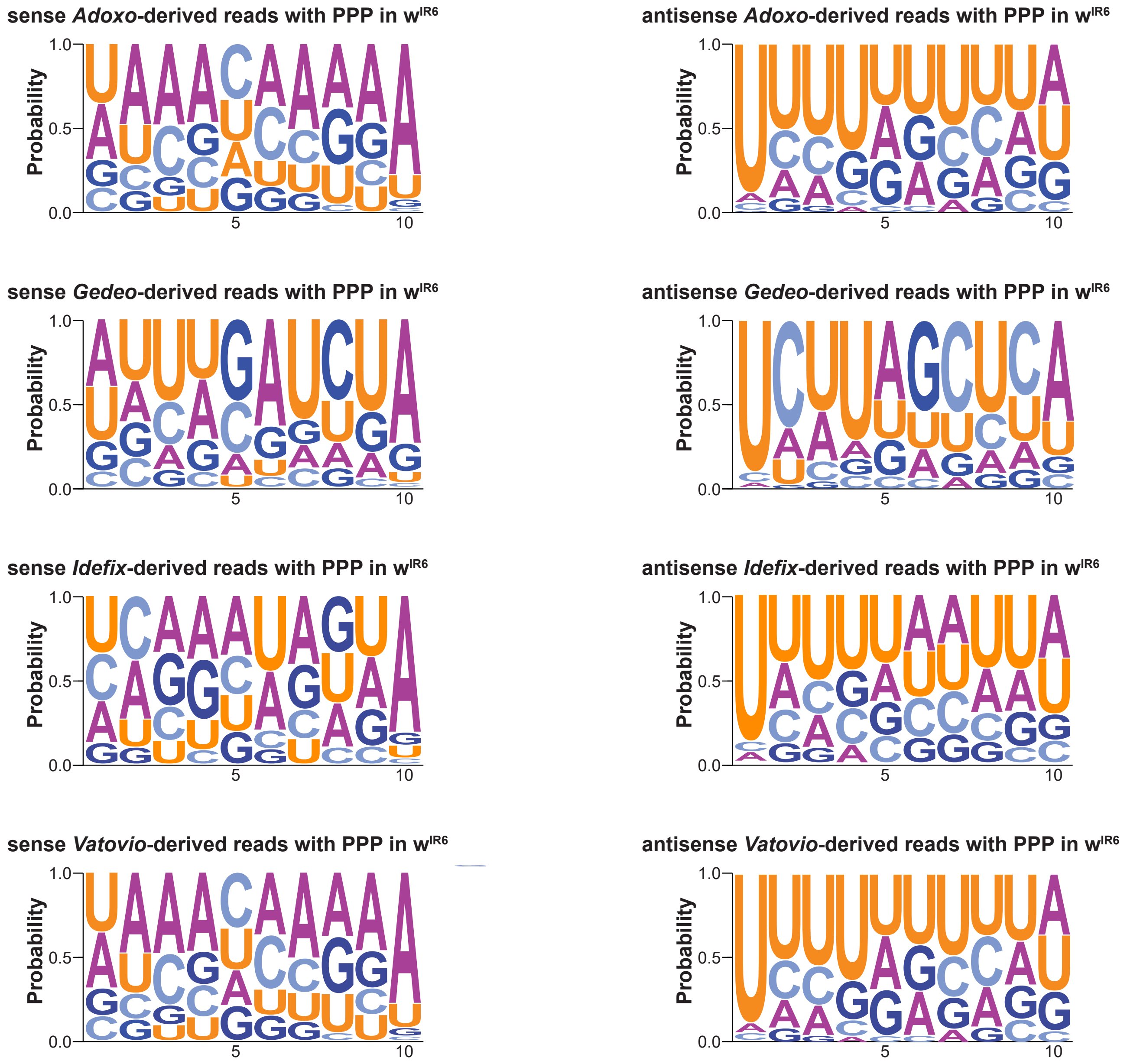

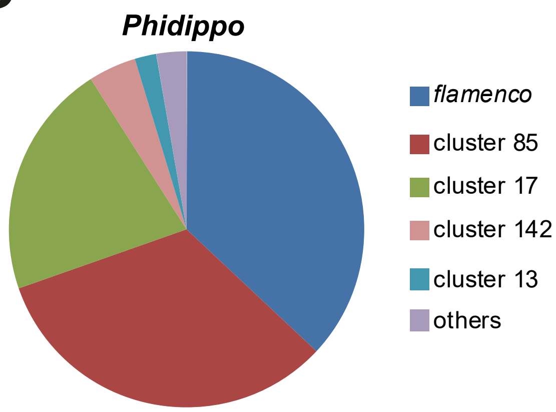

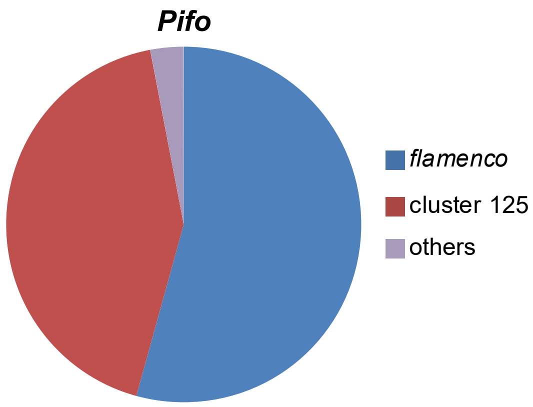

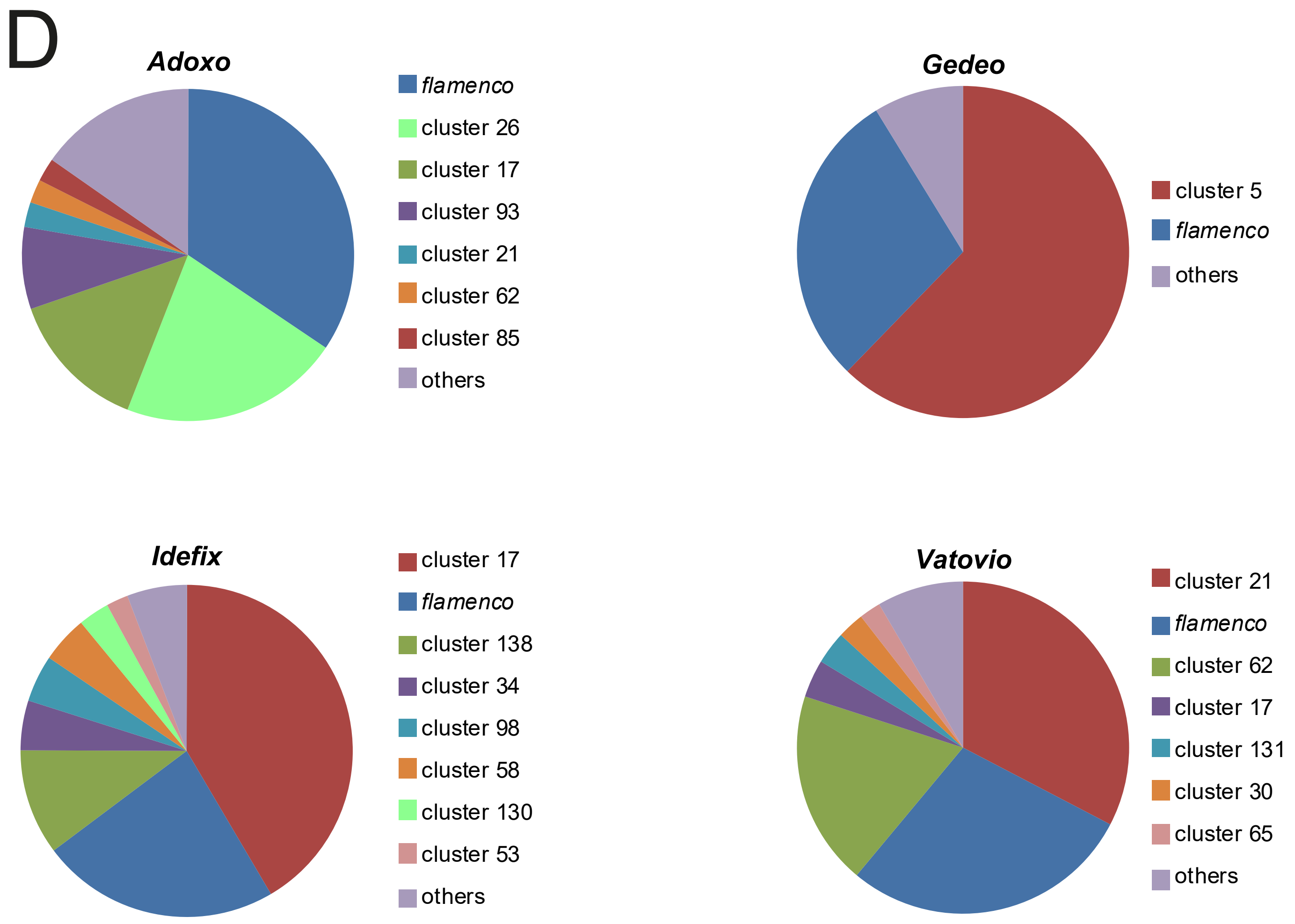

### FigureS6

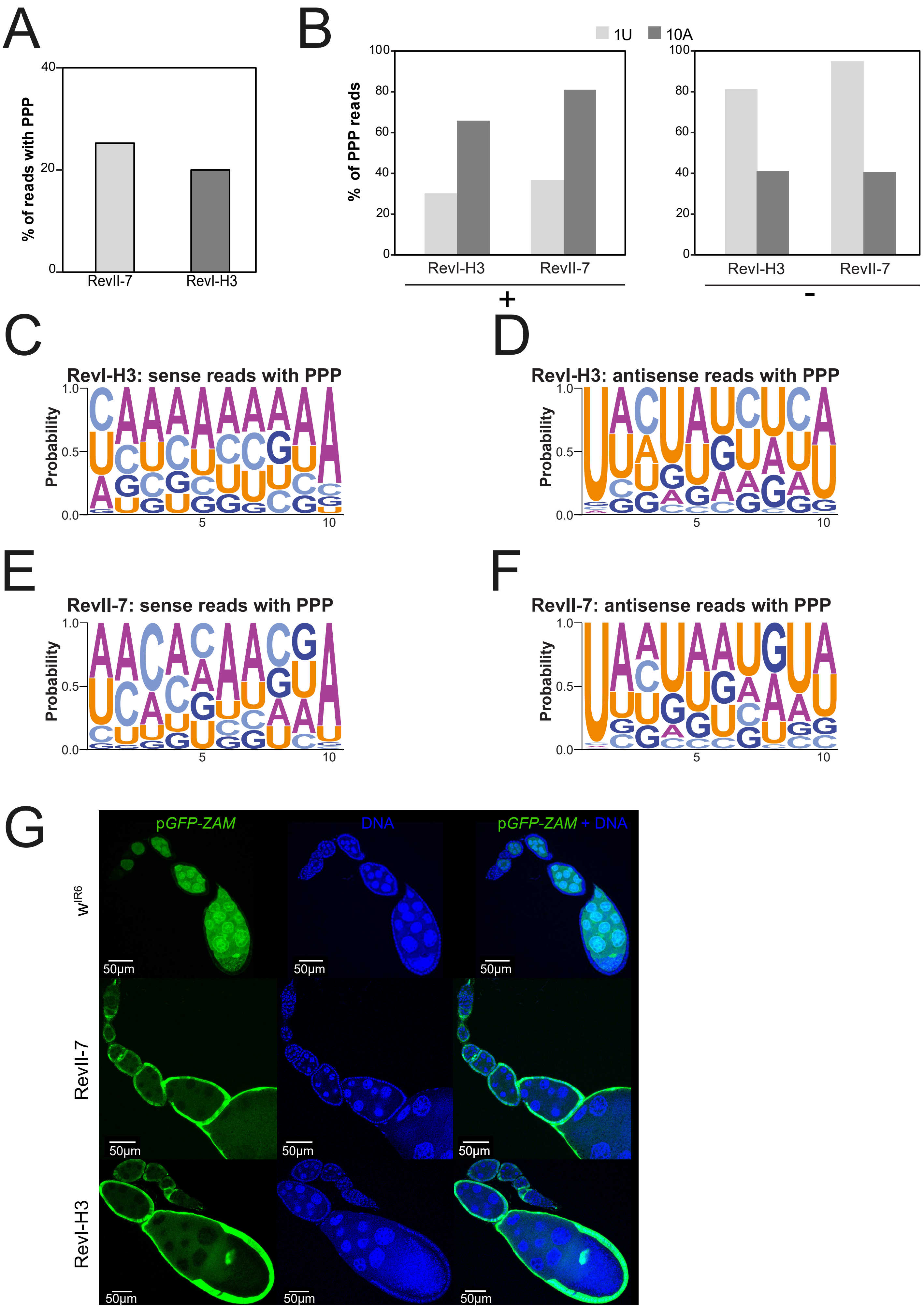

H

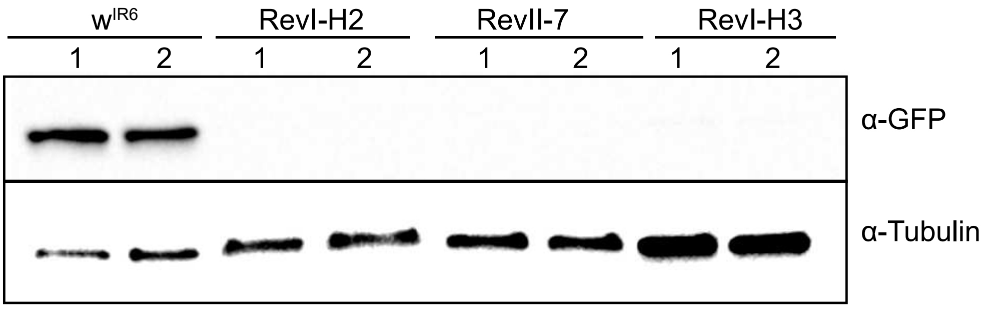
